# *Rhizobium leguminosarum* symbiovar *viciae* strains are natural wheat endophytes and can stimulate root development and colonization by arbuscular mycorrhizal fungi

**DOI:** 10.1101/2020.08.07.241844

**Authors:** Claudia Bartoli, Stéphane Boivin, Marta Marchetti, Carine Gris, Virginie Gasciolli, Mégane Gaston, Marie-Christine Auriac, Ludovic Cottret, Aurélien Carlier, Catherine Masson-Boivin, Marc Lepetit, Benoit Lefebvre

**Affiliations:** IGEPP, INRAE, Institut Agro, Univ Rennes, 35653, Le Rheu, France; LIPM, Université de Toulouse, INRAE, CNRS, 31326 Castanet-Tolosan, France; Laboratoire des Symbioses Tropicales et Méditerranéennes (LSTM) INRAE, IRD, CIRAD, University of Montpellier, Montpellier SupAgro 34398, Montpellier cedex 5, France; FRAIB-TRI Imaging Platform Facilities, Université de Toulouse, CNRS, 24 chemin de Borderouge, 31320 Castanet-Tolosan, France

**Keywords:** *Triticum aestivum* and *Triticum turgidum* (wheat), *Rhizobium leguminosarum* symbiovar *viciae* (*Rlv*), Endophyte, Root development, Arbuscular Mycorrhiza (AM), Competitiveness, Cooperation, Plant Growth-Promoting Rhizobacteria (PGPR)

## Abstract

- Although rhizobia establishing a nitrogen-fixing symbiosis with legumes are also known to promote growth in non-legumes, studies on rhizobia association with wheat roots are scarce.
- We searched for *Rhizobium leguminosarum* symbiovar *viciae* (*Rlv*) strains naturally competent for wheat roots colonization. We isolated 20 strains and tested the ability of a subset for wheat roots colonization when co-inoculated with other *Rlv*. We also measured the effect of these strains on wheat root architecture and Arbuscular Mycorrhizal Fungal (AMF) colonization.
- We found a low diversity of *Rlv* in wheat roots compared to that observed in the *Rlv* species complex. Only a few strains, including those isolated from wheat roots, and one strain isolated from pea nodules, were efficient to colonize wheat roots in co-inoculation conditions. These strains had a high ability for endophytic colonization of wheat root and were able to stimulate root development and AMF colonization in single strain inoculation conditions.
- These results suggest that wheat is an alternative host for some *Rlv*; nevertheless, there is a strong competition between *Rlv* strains for wheat root colonization. Furthermore, our study suggests that the level of endophytic colonization is critical for *Rlv* ability to promote wheat growth.

## Introduction

Plant roots interact with microorganisms that play key roles in their development, nutrition and protection against pathogens (Ortíz-Castro *et al*., 2009). Under the influence of root exudates, microbes multiply in the rhizosphere, i.e. the soil portion in proximity to the roots (Haichar *et al*., 2014: Baetz & Martinoia, 2014; Hu *et al*., 2018; Huang *et al*., 2019). Microorganisms inhabiting the rhizosphere are known to play important roles in plant nutrition through various properties such as biological nitrogen fixation, phosphate solubilization or siderophore secretion (de Souza *et al*., 2015). Among the extremely abundant diversity of soil/rhizospheric microorganisms, only a small fraction is able to colonize the inner part of the plant root system. Several of these endophytic microbes have been shown to offer important benefits to their hosts displaying Plant Growth-Promoting (PGP) activities or protection against pathogens (Santoyo *et al*., 2016). Mechanisms including: i) facilitation in acquiring nutriments, ii) interference with plant hormone (auxin, cytokinin or ethylene) homeostasis, iii) pathogen control via antibiosis (Santoyo *et al*., 2016; Carrión *et al*., 2019) have been related to these beneficial activities.

Among the endophytic microorganisms, some, such as the Arbuscular Mycorrhizal Fungi (AMF) interacting with most of plant species, have the ability to massively colonize plant roots (Bonfante & Genre, 2010). This colonization relies on AMF recognition by plants that is, at least in part, mediated by the perception of fungal LipoChitoOligosaccharide (LCO) signals (Girardin *et al*., 2019). Non-legume LCO receptors are poorly specific for LCO structural variations (Buendia *et al*., 2019a; Girardin *et al*., 2019). Consistently, Arbuscular Mycorrhiza (AM) display poor host-specificity as almost all AMF species are able to colonize most of the terrestrial plants. The LCO perception machinery and the downstream signaling pathway have been recruited in legumes to allow recognition of nitrogen fixing bacteria called rhizobia that are accommodated in root organs called nodules (Girardin *et al*., 2019). Rhizobia also produce LCOs called Nod factors that are recognized by legume hosts (Gough & Cullimore, 2011). Rhizobia are polyphyletic and belong to various bacterial genera. Indeed, the Nod factor synthesis genes (*nod* genes) are frequently located on a plasmid (the symbiotic plasmid), which can be horizontally transferred within or across rhizobia species (Boivin *et al*., 2020a). In contrast with the AM, the rhizobium-legume symbiosis is generally host-specific and symbiovars designate bacteria able to nodulate the same host(s). For example, pea, faba bean, vetch and lentil plants are nodulated by *Rhizobium leguminosarum* bacteria of the symbiovar (sv) *viciae* (*Rlv*), while clover is nodulated by *Rhizobium leguminosarum* sv. *trifolii*. Other legumes such as alfalfa and soybean are nodulated by rhizobia from other genera, *Ensifer* (formerly called *Sinorhizobium*) and *Bradyrhizobium* respectively. Nod factor structural variations are associated to this host-specificity (Gough & Cullimore, 2011). In addition, in natural soils, multiple rhizobia of the same symbiovar coexist but display contrasted Competitiveness to Form Nodules (CFN) depending on their host genotype (Boivin et al, 2020a; Boivin *et al*., 2020b). Together with the host-specificity associated with Nod factor specificity, CFN contribute to determine partner choices during the rhizobium-legume symbiosis.

Several studies have shown that rhizobia can also interact with non-legumes. *Rlv* were previously isolated from soils under wheat monoculture (Depret *et al*., 2004) suggesting that *Rlv* can be maintained in absence of rotation with compatible legumes. Rhizobia belonging to various genera were isolated and/or shown to have PGP activities in rice (Biswas *et al*., 2000; Chaintreuil *et al*., 2000; Peng *et al*., 2002; Yanni & Dazzo, 2010). *R*. *leguminosarum* sv. *trifolii* (isolated from legumes, rice or wheat) were also shown to colonize and have PGP activities on wheat (Höflich *et al*., 1995; Webster *et al*., 1997; Hilali *et al*., 2001; Yanni *et al*., 2016). Although mechanisms involved in AM and rhizobial symbiosis establishment share similarities, there is little information on the rhizobia-AMF interaction, particularly in wheat. Nevertheless, recently Raklami and coworkers showed that rhizobia inoculation in open fields increase wheat colonization by AMF and yield (Raklami *et al*., 2019).

Here we performed a functional ecology study on the wheat-*Rlv* interaction. We investigated whether: i) wheat is a natural host for *Rlv* strains, by isolating bacteria from wheat grown in open fields in the southwest of France ii) there is partner choice in the wheat-*Rlv* interaction, by co-inoculating strains representative of the known *Rlv* diversity iii) there is diversity in *Rlv* for colonization and stimulation of responses in wheat roots, by measuring *Rlv* epiphytic/endophytic colonization, root development and AMF colonization.

## Material and Methods

### Wheat sampling from open fields

Wheat plants (*Triticum aestivum ssp*. *aestivum* and *Triticum turgidum ssp*. *turgidum*) were collected in the CREABio experimental station de la Hourre in Auch (southwest of France; (43°37’17.7”N 0°34’20.6”E) in February 2014. Plants were collected in 2 experimental field plots: LH1 under rotation with pea (*Pisum sativum*) and fertilized with 80 kg/ha of organic fertilizer and LH7 under rotation with soybean (*Glycine max*) and fertilized with 100 kg/ha of organic fertilizer. Both plots were characterized by a limestone clay soil with a pH of 8.1 and 8.4 for LH7 and LH1 respectively. Specific soil composition for the field plots are reported in Fig. S1. Sixteen wheat varieties, 8 from LH1 and 8 from the LH7, were collected (Table S1). Six plants per variety were sampled and transported in sterilized plastic bags.

### Strain isolation and characterization

To isolate *Rhizobium leguminosarum* symbiovar (sv) *viciae* (*Rlv*) strains, 2 methods were employed: nodule-trapping method and direct root plating. Prior to bacterial isolation, wheat roots of each variety were pooled and surface-sterilized. Roots were first rinsed in sterile water to remove soil, and incubated 10 min in a 1.2 % sodium hypochlorite solution, 10 min in a spiramycine 30 mg/ml solution and 1 min in 70% ethanol, followed by 3 rinses in sterile water. Surface-sterilized roots were mixed in 5 ml of phi liquid medium (10 g/L bacto peptone, 1 g/L Casimino acids and 1 g/L yeast extract) and root mixtures were stored in 50% of glycerol at −80°C prior to bacterial isolation. For the nodule-trapping experiment pea (*Pisum sativum*) and vetch (*Vicia sativa*) plantlets were used as specific hosts. Seeds were sterilized and germinated as described in Methods S1. Seedlings were grown in 110 ml glass tubes containing Farhäeus agar medium (Catoira *et al*., 2000). After 10 days, plantlets were inoculated with wheat root homogenates and incubated for 4 weeks at 22°C. For each of the 16 wheat root homogenates, 3 plants of each legume host were inoculated. After 4 weeks, plants were scored for the presence/absence of nodules (Table S1). Each nodule was collected with a scalpel, homogenized and stored in 50% glycerol at −80°C prior to bacterial isolation. Serial dilutions of nodule homogenates were spread on TY agar medium (16 g/L peptone, 10 g/L yeast extract and 5 g/L NaCl) supplemented with 2 mM CaCl_2_. For direct isolation of culturable bacteria, root homogenates were diluted and spread on TY agar medium supplemented with 2 mM CaCl_2._ All purified colonies were first stored in 30% of glycerol/TY medium at −80°C. Bacteria isolated from nodules or by direct plating were tested for presence of the *nodD* gene. We used the Y5 and Y6 primer set developed by Zeze *et al*., (2001) amplifying a 850bp *nodD* fragment from *Rlv, Rhizobium leguminosarum* sv. *trifolii* and *Ensifer meliloti*. Strains showing an amplification were then used for *gyrB* amplification by using primers and protocols already described (Martens *et al*., 2008). Sequences (Table S2) were trimmed and aligned with DAMBE version 6 (Xia, 2017) and neighbor-joining phylogeny in comparison with other *Rhizobium* species (sequences retrieved from GenBank https://www.ncbi.nlm.nih.gov/genbank/; Table S3), was inferred with MEGA 5 (Tamura *et al*., 2011) with *p*-distance model. Three strains isolated from wheat roots, FWPou15, FWPou32 and FW Pou38 were selected for further analysis and their genomes were sequenced and analyzed as described in Methods S2. Genomes are available in GenBank under the accession numbers JACBGR000000000, JACBGQ000000000 and JACBGP000000000, respectively. Phenotypic assays were performed on the 3 strains in comparison with the A34 strain used as control (Götz *et al*., 1985; Oono & Denison, 2010).

### Metabolic pathways and comparative genomics analyses

A metabolic reconstruction in SBML format (Hucka *et al*., 2003) was built for each strain with the Carveme software (Machado *et al*., 2018), a binary matrix containing the presence/absence information for each metabolic reaction in each strain was created from SBML files using ad-hoc Java programs. A correspondence analysis was performed with the dudi.coa method of the R ade4 package (Dray *et al*., 2015). The correspondence analysis result was then plotted using the scatterD3 R package. Tryptophan metabolism was analyzed with the kbase platform (Artkin *et al*., 2018) and by using the RAST annotation pipeline (Brettin *et al*., 2015) and the Model Seed metabolic reconstruction pipeline (Henry et al., 2010). Orthology search was performed with OrthoFinder with default settings (Emms & Kelly, 2019). Functional category assignments and annotations of orthologous groups were done by selecting a representative of each group and annotating with eggNOG-Mapper (Huerta-Cepas *et al*., 2017).

### Nodulation tests

Three selected strains FWPou15, FWPou32 and FWPou38 and the A34 strain were tested for their ability to nodulate pea, faba bean, vetch and clover plants. For pea and vetch plants, germ-free seedlings (Methods S1) were placed into 110 ml glass tubes filled with attapulgite and watered with 10 mL Farhäeus liquid medium supplemented with 1 mM NH_4_NO_3,_ and an aluminum foil was placed around the bottom of the tube. Seedlings were incubated for one week before inoculation with 2 ml of 10^8^/ml bacterial suspension of each strain (12 plant replicates / plant species). For faba bean and clover plants, germ-free seedlings were transferred in containers (250 ml of volume) filled with a mix of 50% perlite/50% sand and humidified with 300 ml of sterilized water. Seedlings were inoculated by adding 150 ml of 10^7^/ml bacterial suspension and cultivated in a growth chamber (20°C and 16h/8h light/dark period). Nodules were counted 6 weeks after inoculation. Microscopy analysis was performed on 3 randomly selected nodules for each nodulated plant species (Methods S3) to confirm that nodules were colonized by bacterial cells.

### Wheat root colonization in co-inoculation assays

Seven-day old germ-free wheat seedlings (Methods S1) of the Energo variety, individually grown in 110 ml glass tubes filled with 70 ml of Fahraeus agar medium (Fig. S2; Methods S4) were inoculated with FWPou15 strain alone or with bacterial mixtures containing either the FWPou15, FWPou32 or FWPou38 strains plus 22 strains of the *Rlv* core collection representing the known genomic *Rlv* diversity (Table S3; Boivin *et al*., 2020b). Each strain present in the mixture was characterized by a unique 309 bp sequence of the *nodD* coding region and were therefore easily unambiguously identified by DNA metabarcoding. Each modality consisted in an inoculation of 6 plants; 3 independent replicates (3 temporal blocks) were performed. In total 18 plants were inoculated with each bacterial mixtures. Each wheat plantlet was co-inoculated with 2 ml of each mixture containing all bacterial strains at the final OD_600nm_= 0.01 (see Table S4 for initial OD_600nm_ of each strain) and incubated in a growth chamber at 20°C and a light/dark period of 16h/8h. Roots were collected after 7 days. Half of the roots were surface-sterilized using the following successive treatments: 1 min in pure ethanol, sterilized water, 3 min in 1.2% sodium hypochlorite solution, 3 times in sterilized water, 1 min in pure ethanol and 3 times in sterilized water. The other half was directly analyzed without surface sterilization. For each temporal block, the plant roots of each modality were pooled prior to DNA extraction with the DNeasy Plant Mini Kit following the manufacturer’s instructions (Table S4). To characterize the relative abundance of the *Rlv* bacteria in the wheat roots, the 309 bp sequence of the *nodD* coding region was amplified by PCR from the DNA extractions and the PCR products were sequenced by Illumina MiSeq 2×250 bp technology as described in Methods S5. Abundance of each strain was estimated by the number of reads corresponding to its unique *nodD* sequence (Methods S5, Table S5). Hierarchical clustering and heatmaps were built using the pheatmap R package (clustering “maximum”, method “wardD”).

### Wheat root colonization in single strain inoculation assays

To test the colonization level in single-inoculation conditions, experiments were performed as described for co-inoculation assays except that plantlets were inoculated with 2 ml of each bacterial suspension (OD_600nm_= 0.08). The number of independent replicates (temporal blocks) and the total number of plants analyzed for each strain and conditions are reported in Table S6. Roots were excised 7 dpi and weighed (fresh weight). Bacterial isolation from roots was performed by chopping the entire root system using a scalpel and by dilution plating on TY plates with the addition of 2 mM CaCl_2_. To estimate the endophytic growth wheat roots were surface-sterilized as described for the co-inoculation assay. Plates were incubated 3 days at 24°C before colony counting. The number of Colony-Forming Units (cfu) was normalized by the root fresh weight.

### Wheat root colonization by the FWPou15 GFP-expressing strain

Construction of the GFP expressing strain is described in Methods S6 and microscopy analysis is detailed in Methods S3.

### Wheat root development assays

Three-days old germ-free wheat seedlings (Methods S1) were placed on 20×20 cm plates containing Fahraeus agar medium (Fig. S2, Methods S4). Plantlets were grown 4 days prior to inoculation with 1 ml of bacterial suspension (OD_600nm_= 0.08) of each strain. Control plants were inoculated with 1 ml of sterilized water. The number of experiments (temporal blocks) and plant replicates are shown in Table S7. Plants were grown at 20°C with a light/dark period of 16h/8h. Roots were removed from the plates 12 dpi, washed with water to remove traces of agar, and imaged with an Epson Expression 10000XL scanner. Total root length was estimated by using WINRhizo software (Pouleur S, 1995) and the number of lateral roots was counted manually on the scanned roots.

### Mycorrhiza assays

Three-day old germ-free wheat seedlings (Methods S1) were transferred in 50 ml containers filled with attapulgite supplemented with 20 ml of 1/2 modified Long Ashton liquid medium (containing 7.5 µM NaH_2_PO_4_) and 200 *Rhizophagus irregularis* DAOM197198 spores (purchased at Agronutrition, Carbonne, France) were inoculated on each plant. The number of experiments (temporal blocks) and plant replicates are shown in Table S8. After 3 days at 23°C with a 16/8h light/dark regime, 1 ml of each bacterial suspension (OD_600nm_= 0.08) was inoculated and plants were further grown for 21 days. Roots were then washed and permeabilized in a 10% KOH solution for 10 min at 95°C and stained with a 5% ink-vinegar solution for 3 min at 95°C.

### Statistical analysis

All the statistical analyses performed in the manuscript were carried out in the R environment. Scripts used for the analyses and for the construction of the graphs are detailed in Methods S7. Raw data that were used to run the statistical models are available in: i) Table S9 for the relative abundances of the *Rlv* strains in roots during the co-inoculation assays, ii) Table S10 for the numbers of cfu in the root colonization assays, iii) Table S11 for the total root lengths and lateral root numbers obtained in the root development assays, iv) Table S12 for the number of colonization sites in the mycorrhiza assays.

## Results

### *Rhizobium leguminosarum* sv. *viciae* (*Rlv*) strains are associated with wheat roots

In order to determine whether *Rlv* strains naturally interact with wheat, we collected wheat roots from plants under rotation with pea or soybean in open fields located in the southwest of France. We surface-sterilized and macerated them to extract bacterial endophytes. Several wheat varieties belonging to 2 species (*Triticum aestivum ssp*. *aestivum* and *Triticum turgidum ssp*. *turgidum*, Table S1) were compared. *Rlv* were isolated from these root samples by two approaches. First, pea and vetch seedlings, which are known legume *Rlv* hosts, were inoculated with the endophyte bacterial suspensions. The nodules formed were individually collected and bacterial clones occupying these nodule were isolated (“nodule trapping”). Second, surface-sterilized macerates from wheat roots were directly plated. Bacteria collected by both methods were PCR screened by using primers that specifically amplify a fragment of the *nodD* gene from *Rlv*, and a few other rhizobial species (Zeze *et al*., 2001). We succeeded in isolating 20 strains from plants in rotation with pea (Table S1). Phylogenetic analysis based on a portion of the chromosomal *gyrB* gene demonstrated that these 20 strains cluster in 2 closely-related clades of the *R*. *leguminosarum* species complex (Fig. 1a). The first clade includes the *Rlv* strains TOM, FRF1D12, Vaf10 and CCBAU83268 belonging to genospecies gsF-1 (Boivin *et al*., 2020a) and the second clade includes the *Rlv* strain SL16 belonging to the genospecies gsF-2 (Boivin *et al*., 2020a). Three strains (FWPou15, FWPou32 and FWPou38) randomly selected from the most predominant clade, were further characterized. Sequencing a large portion of the *nodD* coding region confirmed that they belong to *R*. *leguminosarum* symbiovar *viciae* (Fig. S3). Phylogeny of the *nodD* gene suggested that the symbiotic plasmid of these strains are closely related to that of the reference *Rlv* strain 3841 and the control strain A34 (Oono & Denison, 2010). The 3 strains (FWPou15, FWPou32 and FWPou38) were found to nodulate pea, faba bean and common vetch but not clover plants (Fig. 1b-1e, Fig. S4-S5), confirming that they belong to symbiovar *viciae*.

**Fig. 1.**
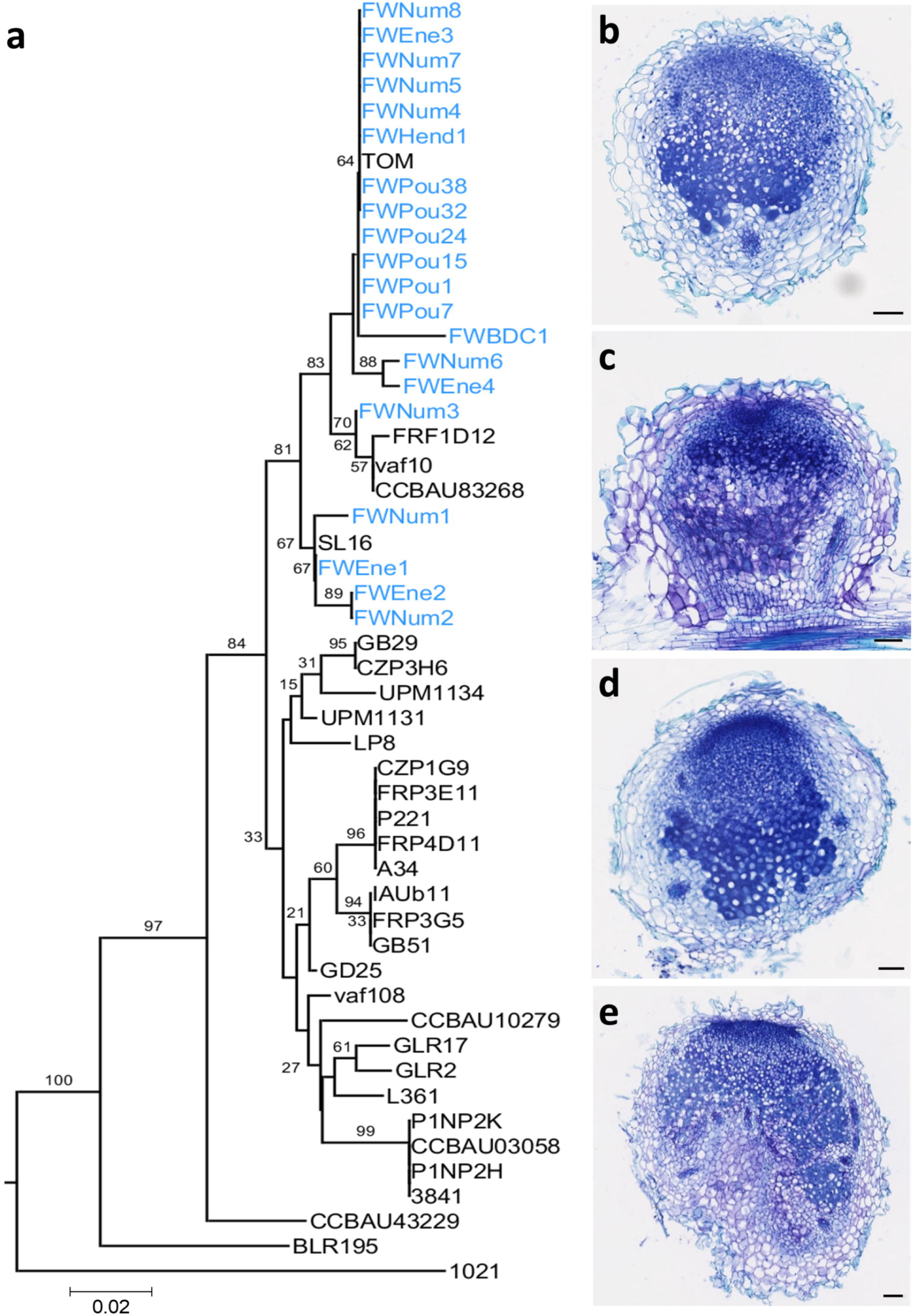
Functional *Rhizobium leguminosarum* sv. *viciae* (*Rlv*) are associated with wheat roots. (a) Neighbor-Joining tree based on a portion of the chromosomic *gyrB* gene from the 20 wheat-isolated strains (indicated in blue), a set of *Rlv* strains representative of the diversity of the species complex (Boivin *et al*., 2020b), and the strain A34. *Ensifer meliloti* was used as outgroup. The *Rlv* strains isolated from wheat roots cluster with strains from genospecies gsF-1 (TOM, FRF1D12, Vaf10, CCBAU83268) or gsF-2 (SL16). (b-g) Sections of pea nodules colonized by (b) FWPou15, (c) FWPou32, (d) FWPou38 and (e) A34. Scale bars correspond to 100 µm.

### Wheat-associated *Rlv* do not display specific genomic and metabolic signatures

Whole genome sequencing was performed on FWPou15, FWPou32 and FWPou38. Genomes of these strains were compared to genomes of *Rlv* isolated from legumes and representative of the gsB, gsC, gsE and gsF-1/2 genospecies (Boivin *et al*., 2020a). Sequences of the 3 strains shared a ca. 100% Average Nucleotide Identity (ANI) and few SNPs were detected between the strains. Genomic phylogeny confirmed that the 3 wheat strains cluster with the representatives of genospecies gsF-1 (Fig. 2). Phylogeny on 11 concatenated *nod* genes located on the symbiotic plasmid, showed that the 3 wheat strains cluster with the Nod group B1 that includes the 3841 and FRF1D12 strains isolated from faba bean and pea respectively (Fig. 3; Boivin *et al*., 2020a). The genomes of strains FWPou15, FWPou32 and FWPou38 encode 301 orthologous genes, which are not shared with other *Rlv* representatives. A majority of these genes (200/301) were not assigned to a specific Clusters of Orthoulogous Groups (COG) functional category. We could not identify functions with an obvious link to symbiotic or endophytic growth among the remaining genes. We further inferred and compared the metabolic potential of the 3 wheat strains and 22 strains isolated from legume nodules, representing the *Rlv* genetic diversity (Table S3, Boivin *et al*., 2020b). We found that all the strains shared most of the metabolic reactions (1,967 reactions, >87%; Fig. S6a) and only 13% of the reactions were specific to a subset of strains. Correspondence Analysis (CA) constructed on the presence/absence of the metabolic reactions, showed a hierarchical clustering similar to that observed for the ANI (Fig. S6b; Fig. 2), in which the wheat isolates group with the TOM strain. In conclusion, wheat-isolated *Rlv* do not display specific genomic and metabolic signatures when compared with *Rlv* isolated from legume nodules.

**Fig. 2.**
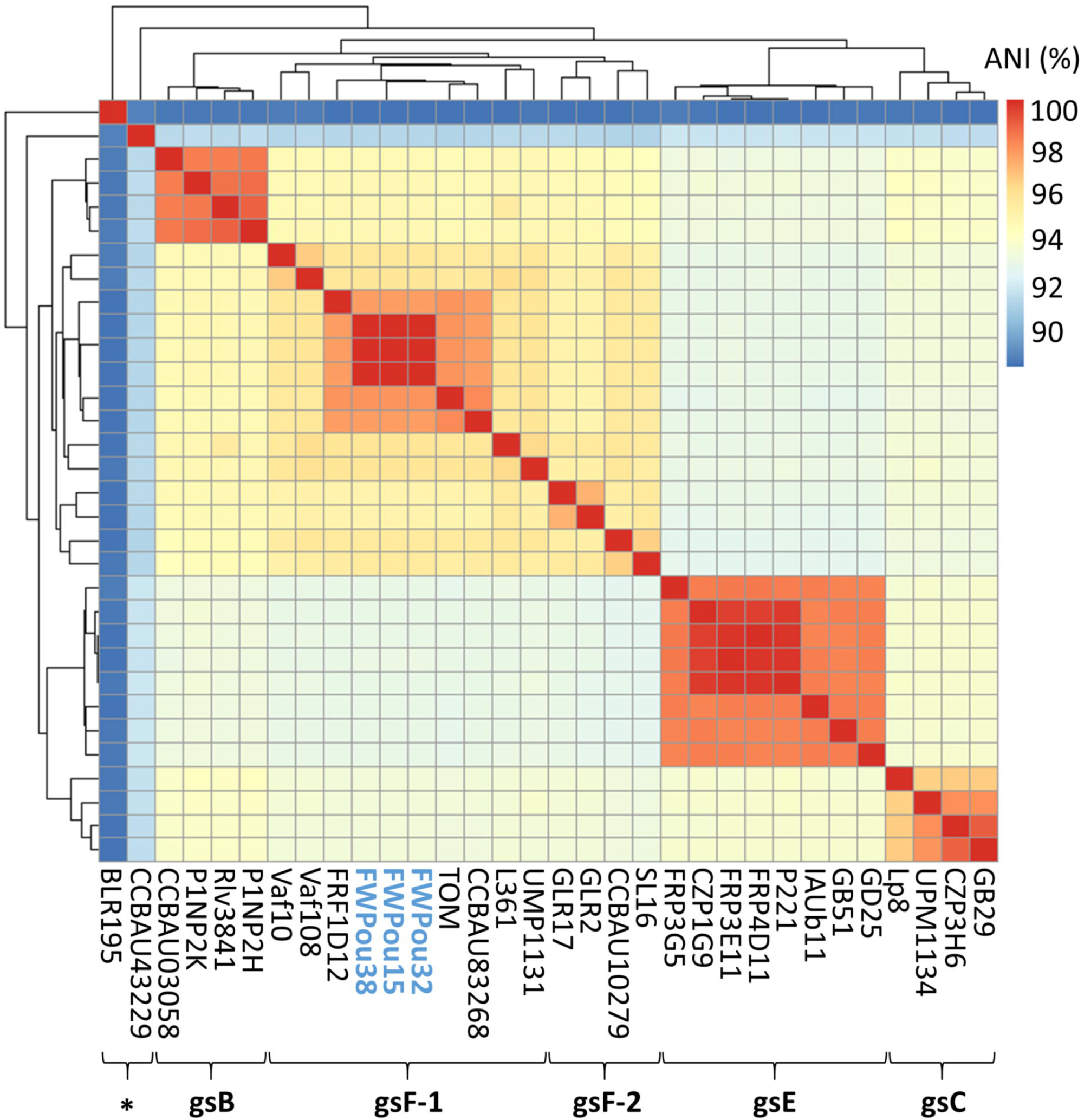
Selected *Rlv* strains for functional analyses belong to genospecies gsF-1. Hierarchical clustering and heat map based on the Average Nucleotide Identity (ANI) values between each couple of the 3 wheat strains, FWPou15, FWPou32 and FWPou38 (indicated in blue), and a set of *Rlv* strains representative of the diversity of the species complex (Boivin *et al*., 2020b). Genospecies (gs) have been defined using an ANI threshold of 95%. Star indicates phylogenetically distant isolates that do not cluster with any of the genospecies.

**Fig. 3.**
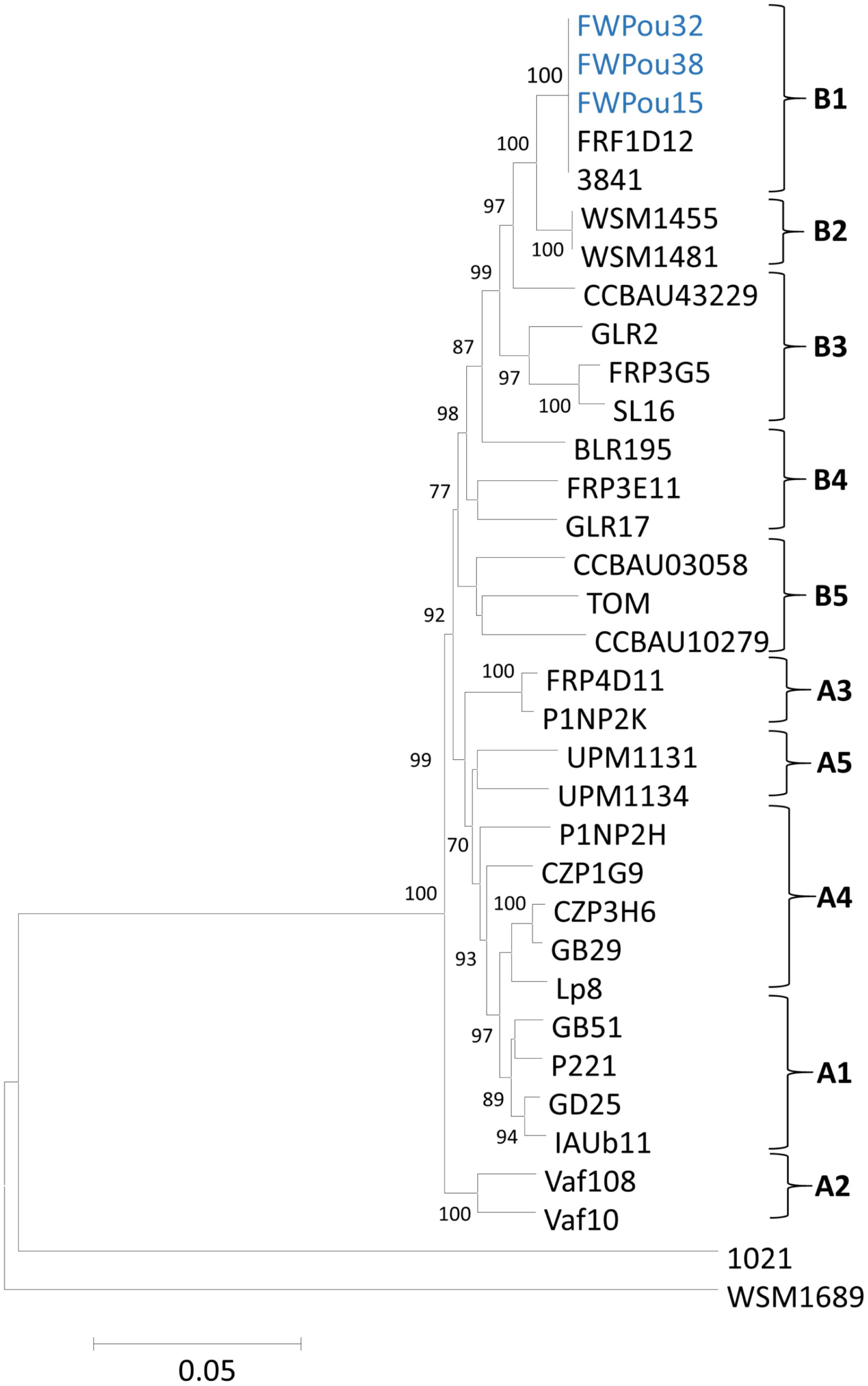
Selected *Rlv* strains for functional analyses belong to the Nod group B1. Neighbor-Joining tree based on concatenated *nodABCDEFIJLMN* gene sequences of the 3 wheat strains, FWPou15, FWPou32 and FWPou38 (indicated in blue), and *Rlv* strains representative of the Nod groups (Boivin et al., 2020b). WSM1689 (*R*. *leguminosarum* symbiovar *trifolii*) and *E*. *meliloti* 1021 were used as outgroups.

### *Rlv* strains colonize wheat roots with different degrees of success in co-inoculation assays

We then tested whether there is diversity among the *Rlv* complex species for wheat root colonization ability. For this, we compared the colonization success of wheat isolates with that of strains isolated from legume nodules in a co-inoculation experiment. We inoculated in gnotobiotic conditions, plantlets of the *T*. *aestivum* Energo variety with 3 mixtures containing an equal concentration of 23 strains, each mixture consisting of the 22 strains representing the genetic diversity of *Rlv* (Table S3) and either FWPou15, FWPou32 or FWPou38. In each replicate, plantlets were also inoculated with FWPou15 alone to verify the efficacy and specificity in detecting the *Rlv* strains. We quantified the relative abundance of each strain 7 days post inoculation (dpi) by Illumina MiSeq sequencing of a *nodD* gene fragment (Methods S5, Boivin *et al*., 2020b) on DNA extracted after surface-sterilization of the roots (treatment S) or without root surface-sterilization (treatment NS). In the control plants inoculated with FWPou15 alone, on average 95% of the total reads corresponded to this bacterium (Fig. S5) confirming its ability to colonize wheat roots and suggesting little contamination in the experiment. In the co-inoculated plants, thirteen strains were either not detected (no reads, CZP3H6, GD25 and CCBAU03058, Table S5) or had a relative abundance lower than 1% (BLR195, CCBAU10279, FRP3E11, GB51, GLR2, GLR17, P1NP2K, SL16, TOM, UPM1134; Fig. 4, Table S5), suggesting there are either not able to colonize wheat roots or poor competitors. The other strains including ten strains isolated from legumes as well as the wheat strains were detected both in NS and S roots, revealing wheat root epiphytic and endophytic colonization abilities are widely distributed among the *Rlv* diversity. However, strong differences in the relative abundance of these strains were observed. A generalized mixed-linear model (*glmm*) on the relative abundances of the strains in the 3 mixtures showed a significant effect of the ‘*strain’* factor, but not of the ‘*sterilization*’ factor, on the colonization success (Table 1). The IAUb11 strain isolated from pea nodules was significantly more abundant than all the other strains in both root compartments (Table S13), with a mean relative abundance of 63% and 45% in NS and S roots, respectively (Fig. 4, Table S5). The wheat strains FWPou32 and FWPou38 showed intermediate colonization abilities, with a mean relative abundance of 13 and 4% in NS roots and 19 and 22 % in S roots (Fig. 4, Table S5). Significant differences in their abundance compared to other strains (except IAUb11) were only found in the S roots (Table S13). Interestingly, the strain TOM, belonging to the same gsF-1 genospecies as the wheat strains, had very low ability to colonize wheat roots in this assay (read abundance <1%) suggesting that the genospecies does not predict wheat colonization ability.

**Table 1.** Analysis of Deviance on the variation of the strain relative abundance inferred from the total number of reads obtained by Illumina MiSeq sequencing of a *nodD* fragment in the co-inoculation assays. Chisq: value of the type II Wald chi squared, Df: degree of freedoms. Significant results obtained after FDR correction are reported in bold.

| FWPou15 mix |  |  |  | FWPou32 mix |  |  | FWPou38 mix |  |  |
| --- | --- | --- | --- | --- | --- | --- | --- | --- | --- |
| Factors | Chisq | Df | P value | Chisq | Df | P value | Chisq | Df | P value |
| Sterilization | 0.08 | 1 | 0.7775 | 0.0012 | 1 | 0.945 | 0.0012 | 1 | 0.9727 |
| Strain | 545.25 | 19 | **<2e-16** | 482.07 | 19 | **<2e-16** | 605.93 | 19 | **<2e-16** |
| Treatment : Strain | 27.05 | 19 | 0.1035 | 26.57 | 19 | 0.1151 | 24.83 | 19 | 0.1662 |

**Fig. 4.**
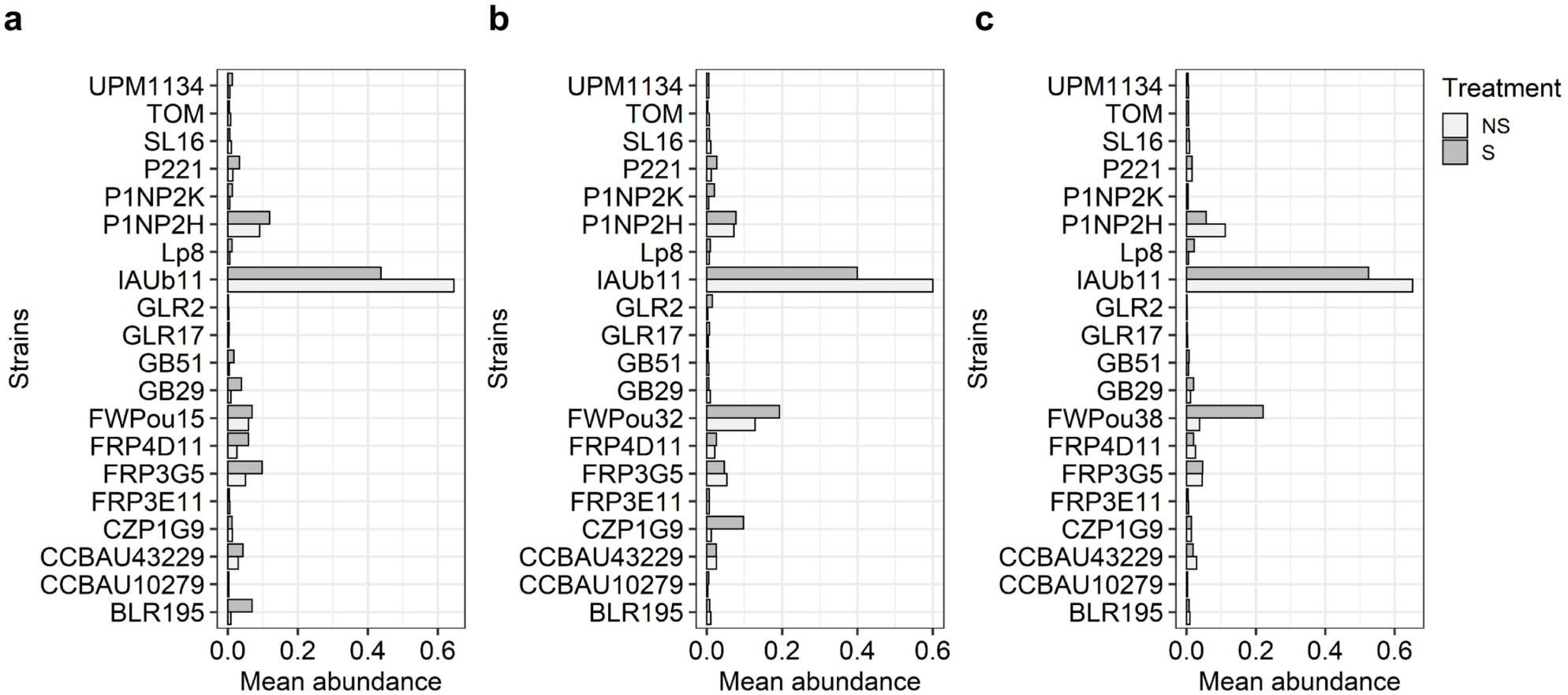
*Rlv* strains colonize wheat roots with varying degrees of success in co-inoculation assays. (a-c) Least-Squares-Means (lsmeans) of the *Rlv* strain abundance in wheat roots when co-inoculated in an *in vitro* assay. Co-inoculation of 22 *Rlv* strains representing the diversity of *Rlv* (Table S3) together with the wheat strains FWPou15 (a), FWPou32 (b) or FWPou38 (c). Each mixture was inoculated in a total of 18 wheat plant root systems in 3 independent replicates. Total DNA was extracted from pools of 3 roots systems, with or without surface-sterilization, at 7 dpi, and a *nodD* amplicon was sequenced by MiSeq Illumina. Mean abundance was inferred by normalizing the number of reads corresponding to each strain with the total number of reads obtained in the run. Dark grey bars indicate results obtained from surface-sterilized (S) roots and light grey bars indicate results obtained from non-sterilized (NS) roots. The S and NS treatments allow estimating respectively the relative endophytic and epiphytic abundance of each inoculated strain since endophytic bacteria are in negligible amounts compared to epiphytic bacteria. Three *Rlv* strains GD25, CCBAU03058 and CZP3H6 were not detected by MiSeq sequencing, thus, they are not reported in the barplots.

The colonization success of each *Rlv* strain in the mixtures is globally equivalent in S and NS roots, suggesting that epiphytic colonization success drives the endophytic colonization success. However, the strains FWPou32 and particularly FWPou38 were more successful in colonizing S roots than NS roots, suggesting a stronger endophytic ability compared to most of the *Rlv* strains isolated from legumes.

### *Rlv* strains are wheat root endophytes

We then quantified the endophytic and epiphytic wheat root colonization abilities of a few individual strains. We selected the strains IAUb11, FWPou38 and FWPou15 displaying respectively the high, intermediate and low degree of colonization in co-inoculation conditions. We included in the analysis the commonly used *Rlv* strain A34 as a control. We performed single strain inoculations in gnotobiotic conditions on two *T*. *aestivum* varieties, Energo and Numeric, and colonization was assessed by measuring cfu of strains isolated from wheat roots 7 dpi and following surface-sterilization (treatment S) or not (NS) of the wheat root system.

A linear-mixed model (*lmm*) on the cfu /mg of root fresh weight showed a significant effect of the ‘*sterilization*’ and ‘*strains*’ factors (Table 2). No significant effect was observed for the ‘*variety*’ factor, however, significant nested ‘*sterilization* × *variety*’ factor and ‘*strain* × *variety*’ effects were observed (Table 2).

**Table 2.**
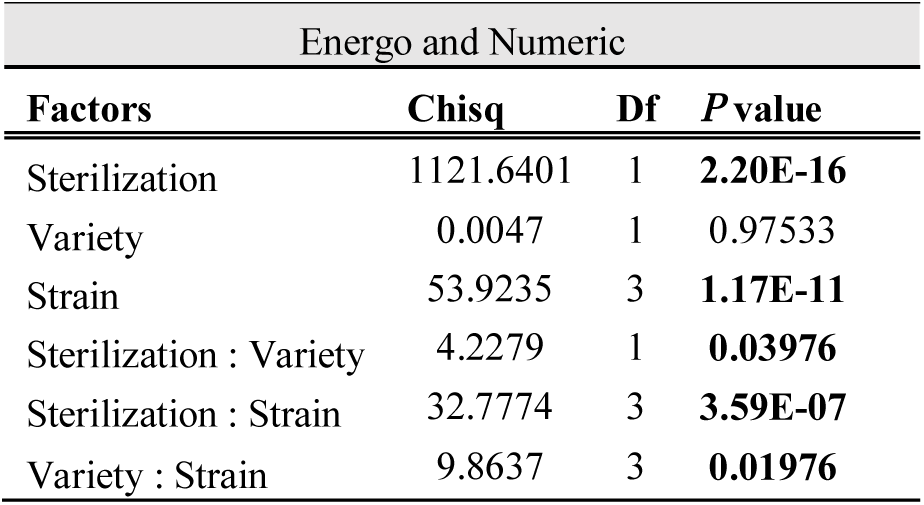
Analysis of Deviance on the variation of bacterial colony-forming units (cfu) / mg of wheat roots obtained after inoculation with the A34, FWPou15, FWPou38 or IAUb11 *Rlv* strains. Chisq: value of the type II Wald chi squared, Df: degree of freedoms. Significant results obtained after FDR correction are reported in bold.

All strains colonized the root surface of the 2 wheat varieties (10^6^ to 10^7^ cfu/mg, Fig. 5) and no significant difference was observed between strains and between the wheat varieties in the NS roots (Table S14). By contrast, bacteria differed in their endophytic colonization abilities. FWPou38 and IAUb11 were the most efficient in both varieties (10^3^ to 10^4^ cfu/mg), while A34 was the least efficient root endophyte (10^2^ cfu/mg, Fig.5) on Numeric. FWPou15, which behaved similarly to A34 on the Energo variety, displayed an intermediate endophytic colonization ability on Numeric (10^3^ cfu/mg, Fig. 5, Table S14).

**Fig. 5.**
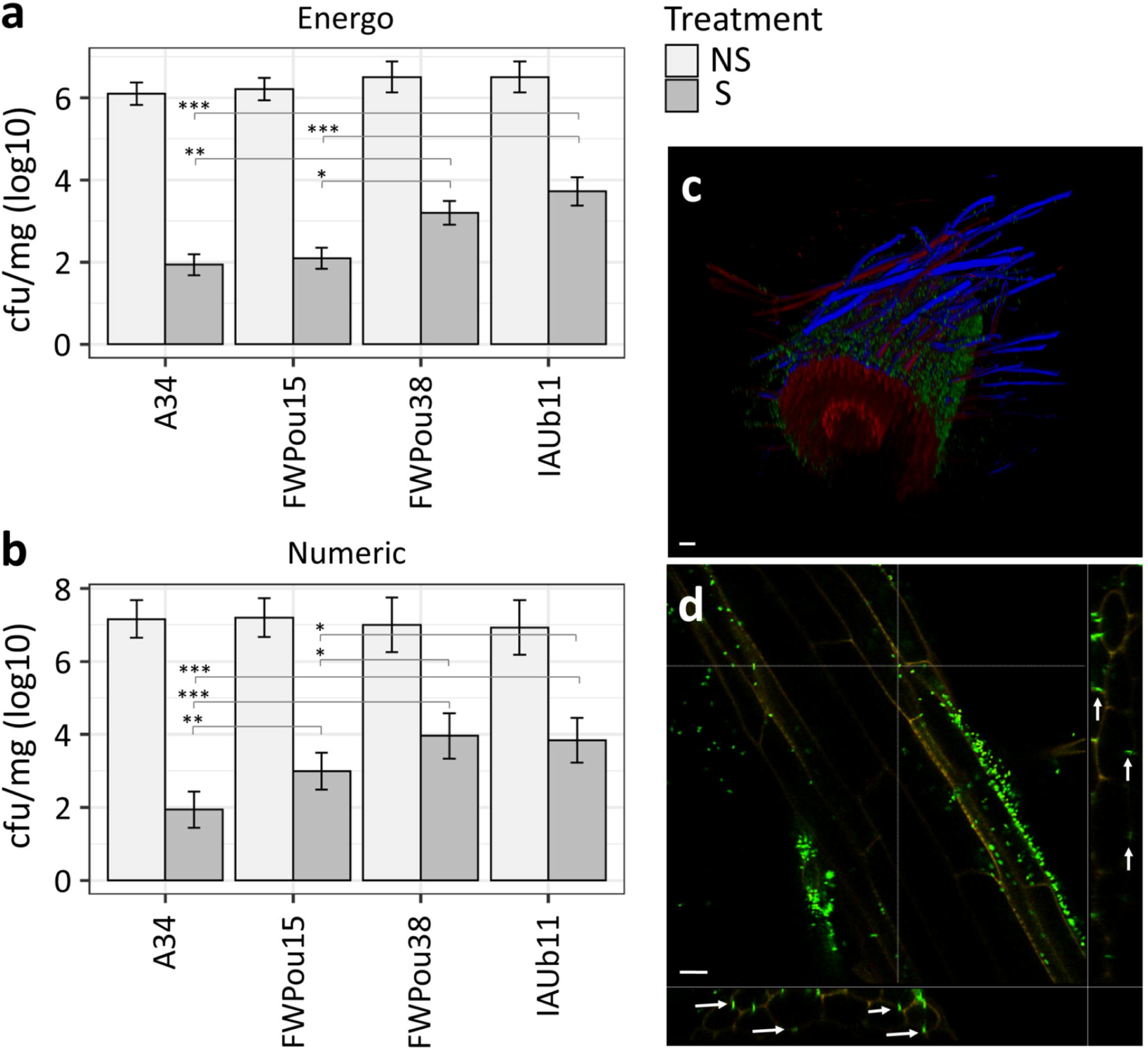
*Rlv* strains are wheat root endophytes. (a-b) Barplots representing the endophytic (surface-sterilized: S; dark grey bars) and epiphytic (non-sterilized: NS; light grey bars) bacterial abundance in roots of the wheat varieties Energo (a) or Numeric (b). Strain name is reported in the *x*-axis. Least-Squares-Means (lsmeans) and standard deviation of bacterial abundance 7 dpi is expressed as log_10_ of cfu (colony-forming units)/mg of root fresh weight. Total number of plants analyzed (≥ 6 for NS treatment and 12 for S treatment, in at least 2 independent replicates) are indicated in Table S6. Significant differences were found in the surface-sterilized roots. *P* values (corrected for FDR) ≤ 0.05 are indicated by *, ≤ 0.005 by ** and ≤ 0.0005 by *** (See Table S14). (c-d) Confocal microscope images of Energo wheat roots inoculated with the FWPou15 GFP-tagged strain. 3D reconstruction of a cleared root fragment (c). Root median focal plan (central) and transversal section reconstructions (bottom and right) of a root fragment in absence of clearing (d). Green fluorescence corresponds to FWPou15-GFP. Red and blue fluorescence correspond to root cell wall auto fluorescence. Arrows indicate the presence of endophytic bacteria between epidermal and cortical cells. Scale bars corresponds to 20 µm.

To confirm the epiphytic and the endophytic capacities of *Rlv* through microscopy observation, we constructed a GFP-tagged version of FWPou15 and observed its *in vitro* root colonization on the cultivar Energo 7 dpi. Confocal microscopy showed that FWPou15 massively colonized the surface of wheat roots (Fig. 5c). A few bacteria were also observed inside the roots, at least around the outer cortical cells (Fig. 5d), confirming their endophytic colonization ability.

### *Rlv* can stimulate wheat root development

We then tested whether *Rlv* strains show differences in their ability to induce plant responses. We first investigated whether *Rlv* strains have different effects on root architecture. We inoculated A34, FWPou15, FWPou38 or IAUb11 on both Energo and Numeric varieties in gnotobiotic conditions, measured the number of lateral roots and the total root length at 12 dpi. *lmm* on both variables showed significant effects of the ‘*strain’* and ‘*variety*’ factors as well as a significant nested ‘*strain* x *variety*’ effect (Table 3). Despite that for each wheat variety the total root lengths and the lateral root numbers were correlated (*R*^2^ ∼0.7, Fig. S7), these traits responded differently to inoculation (Fig. 6). FWPou38 and IAUb11 induced a significant increase in the lateral root number (on average + 35% each; Fig. 6d, Table S15), while FWPou15 induced a weak but significant effect on the total root length (on average +12 %; Fig. 6a, Table S15) in the Energo variety. In contrast, FWPou15 induced a significant increase of both total root length (on average +29%; Fig. 6a, Table S15) and lateral root number (on average +52%; Fig. 6b, Table S15) in the Numeric variety; while FWPou38 induced a significant, although weak, decrease in the total root length (on average -18 %; Fig. 6d, Table S11). Inoculation by A34 did not affect root architecture in either wheat variety. Interestingly, on the Energo variety, strains with the highest level of endophytic colonization (FWPou38 and IAUb11) showed the strongest effect on wheat root architecture. FWPou15 displayed a root-growth activity mostly on the Numeric variety in which it showed a higher level of endophytic colonization compared to the Energo variety (Fig. 4).

**Table 3.**
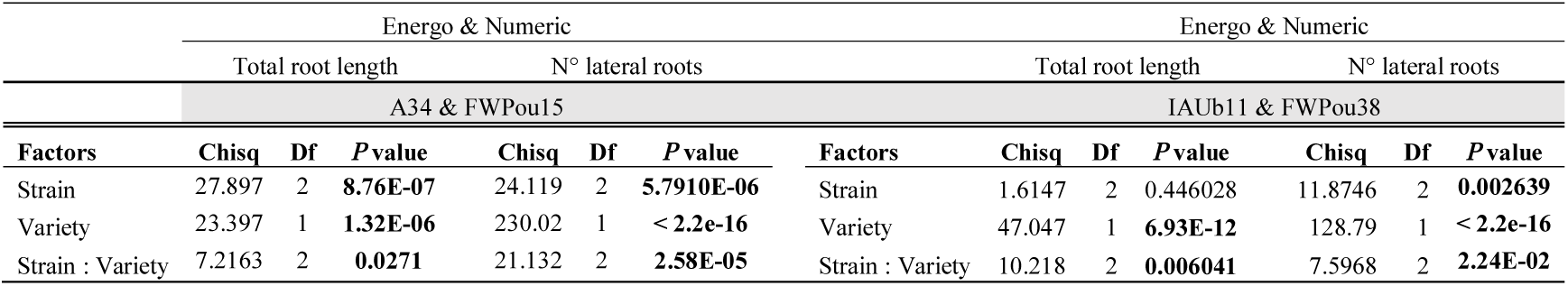
Analysis of Deviance on the variation of the total root length and number of lateral roots obtained after inoculation with the A34, FWPou15, FWPou38 or IAUb11 *Rlv* strains. Analysis was performed on two datasets: A34-FWPou15 and IAUb11-FWPou38, as values for controls (plantlets inoculated with water) were different in the two datasets. Chisq: value of the type II Wald chi squared, Df: degree of freedoms. Significant results obtained after FDR correction are reported in bold.

**Fig. 6.**
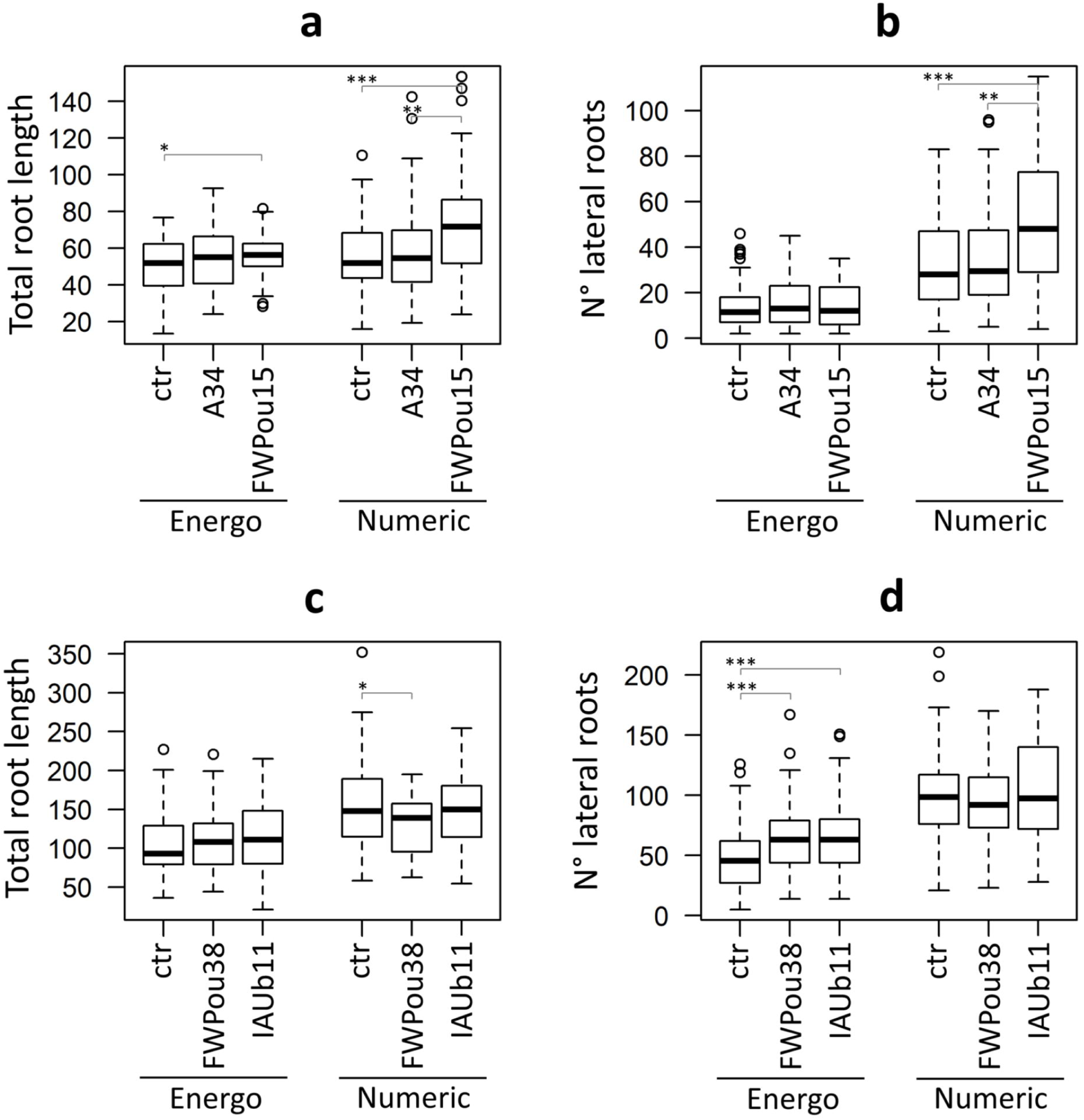
*Rlv* can stimulate wheat root development in a genotype-dependent manner. Boxplots representing the variation in wheat total root length (a, c; cm) and lateral root number (b, d) 12 dpi in 2 wheat varieties (Energo and Numeric) inoculated with the strains A34 or FWPou15 (a, b), FWPou38 or IAUb11 (c, d). Controls (ctr) are non-inoculated plants. Total numbers of plants analyzed (≥ 30, in at least 2 independent replicates) are indicated in Table S7. *P* values (corrected for FDR) ≤ 0.05 are indicated by *, ≤ 0.005 by ** and ≤ 0.0005 by *** (See Table S15).

### *Rlv* can enhance colonization of wheat roots by arbuscular mycorrhizal fungi

We then investigated whether *Rlv* strains have different effects on wheat root colonization by AMF. For this, we co-inoculated Energo and Numeric roots with the *Rhizophagus irregularis* isolate DAOM197198 and one of the 4 *Rlv* strains (FWPou15, FWPou38, IAUb11 or A34). We measured the number of AMF colonization sites 24 dpi and compared it to wheat plantlets inoculated only with *R*. *irregularis*.

*lmm* showed a significant effect of the ‘*strain*’ factor only on the Energo variety (Table 4). More specifically, when compared to the control, inoculation with the FWPou38 and IAUb11 strains resulted in significantly more fungal colonization sites (on average +55% and +31% respectively; Fig. 7b; Table S16). Similarly, roots inoculated with either FWPou38 or IAUb11 displayed significantly more fungal colonization sites than roots inoculated with A34 (on average +77% and +50% respectively; Table S16). Only FWPou38 induced significantly more fungal colonization sites than FWPou15 (on average +33%; Table S16). On the Numeric variety, inoculation with any of *Rlv* strain did not result in a significant change in the number of fungal colonization sites (Fig. 7c and d; Table S16) reinforcing the influence of the wheat genotype on the response induced by *Rlv*. Taken together, these results suggest that the *Rlv* strains stimulating AMF colonization in wheat roots are those able to stimulate the lateral root number.

**Table 4.**
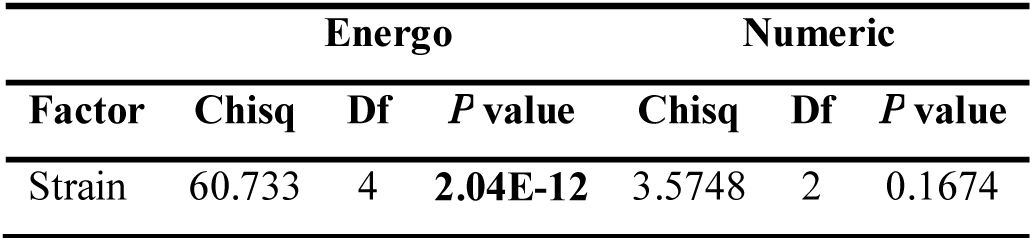
Analysis of Deviance on the variation of the number of AMF colonization sites after inoculation with the A34, FWPou15, FWPou38 or IAUb11 *Rlv* strains. Chisq: value of the type II Wald chi squared; Df: degrees of freedom. Strain refers to wheat roots inoculated with the four *Rlv* strains or sterilized water (control). Significant result is reported in bold corresponding to P-value after FDR correction.

| Factor | Energo |  |  | Numeric |  |  |
| --- | --- | --- | --- | --- | --- | --- |
|  | Chisq | Df | *P* value | Chisq | Df | *P* value |
| Strain | 60.733 | 4 | **2.04E-12** | 3.5748 | 2 | 0.1674 |

**Fig. 7.**
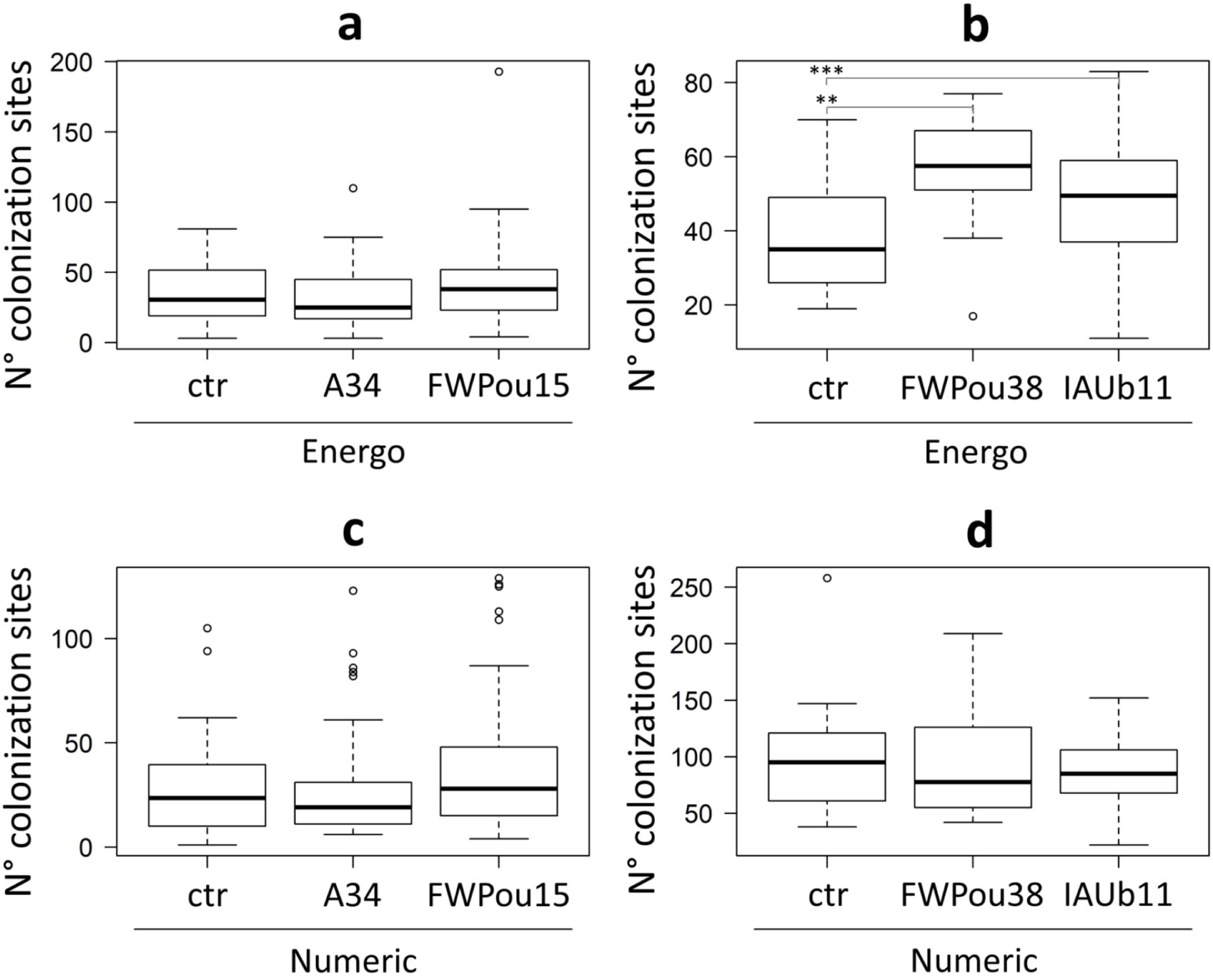
*Rlv* can stimulate wheat root colonization by AMF in a genotype-dependent manner. (a-c) Boxplots representing the variability in the number of colonization sites by the AMF *R*. *irregularis* isolate DAOM197198, 24 dpi, when inoculated alone (ctr) or in combination with *Rlv* strains. Two wheat varieties, Energo (a, b) and Numeric (c, d) were co-inoculated with the strains A34 or FWPou15 (a, c), FWPou38 or IAUb11 (b, d). Total numbers of plants analyzed (≥ 31, in at least 2 independent replicates) are indicated in Table S8. *P* values (corrected for FDR) ≤ 0.005 are indicated by ** and ≤ 0.0005 by *** (See Table S16). Significant differences, not represented on the boxplots, were also found in the Energo variety between A34 *vs* FWPou38 (*P* **<**.0001) or IAUb11 (*P* = 0.0007), and FWPou15 *vs* FWPou38 (*P* = 0.0062).

## Discussion

### Wheat might participate in shaping *Rlv* populations in agronomical soils

*Rhizobium leguminosarum* sv. *viciae* strains are known to nodulate legume hosts such as pea and faba bean and have PGP activities on wheat. Here we show that wheat is an alternative natural host for *Rlv*. Yet, in our study we only isolated *Rlv* strains from root samples of wheat plants cultivated in rotation with pea plants (*Rlv* host), while we did not succeed in isolating *Rlv* from wheat plants in rotation with soybean (non-*Rlv* host). This suggests an enrichment of *Rlv* in wheat roots through rotation with a legume host. On the other hand, a previous study described *Rlv* strains in soils under wheat monoculture (Depret *et al*., 2004). Although the *Rlv* species complex had been currently described to contain 5 genospecies, only members of two of them (gsF-1 and gsF-2) were identified among the 20 wheat isolated strains. The limited diversity of *Rlv* found in our wheat samples might be explained through three non-exclusive hypotheses. Firstly, the sampling we performed was not saturated. Secondly, we cannot exclude that soils in which wheat have been grown were deprived of other *Rlv* genospecies. However, this is not in accordance with previous results showing a high *Rlv* diversity in European soils including in the southwest of France (Boivin *et al*., 2020a). Thirdly, the low *Rlv* diversity is related to partner choice in wheat-*Rlv* interactions. Nevertheless, through co-inoculation assays, we demonstrated that *Rlv* strains isolated from legume plants and belonging to other genospecies (i.e. IAUb11, gsE) can efficiently colonize wheat roots while other members of gsF-1/2 genospecies were not efficient wheat root colonizers suggesting that wheat partner choice is not related to *Rlv* genospecies. However, the co-inoculation assays resulted in a reduced diversity associated with the wheat roots compared to the inoculum, supporting the hypothesis of partner choice. Such partner choice was also observed in the legume-*Rlv* interaction (Boivin *et al*., 2020b). Interestingly, the pattern of colonization success we found for the *Rlv* strains in the wheat roots was different to that observed with the same 22 *Rlv* for colonization in various legumes during nodulation (Fig. 8; Boivin *et al*., 2020b), suggesting that *Rlv* strains might carry different competitiveness abilities to colonize legume nodules, wheat roots or other putative hosts. By consequence, multiplicity of plant hosts in fields can participate in maintaining *Rlv* diversity in soils. Field campaigns at a broader geographical scale to investigate the diversity of *Rlv* in wheat roots, legume nodules and in the surrounding soils, should help to validate these hypotheses.

**Fig. 8.**
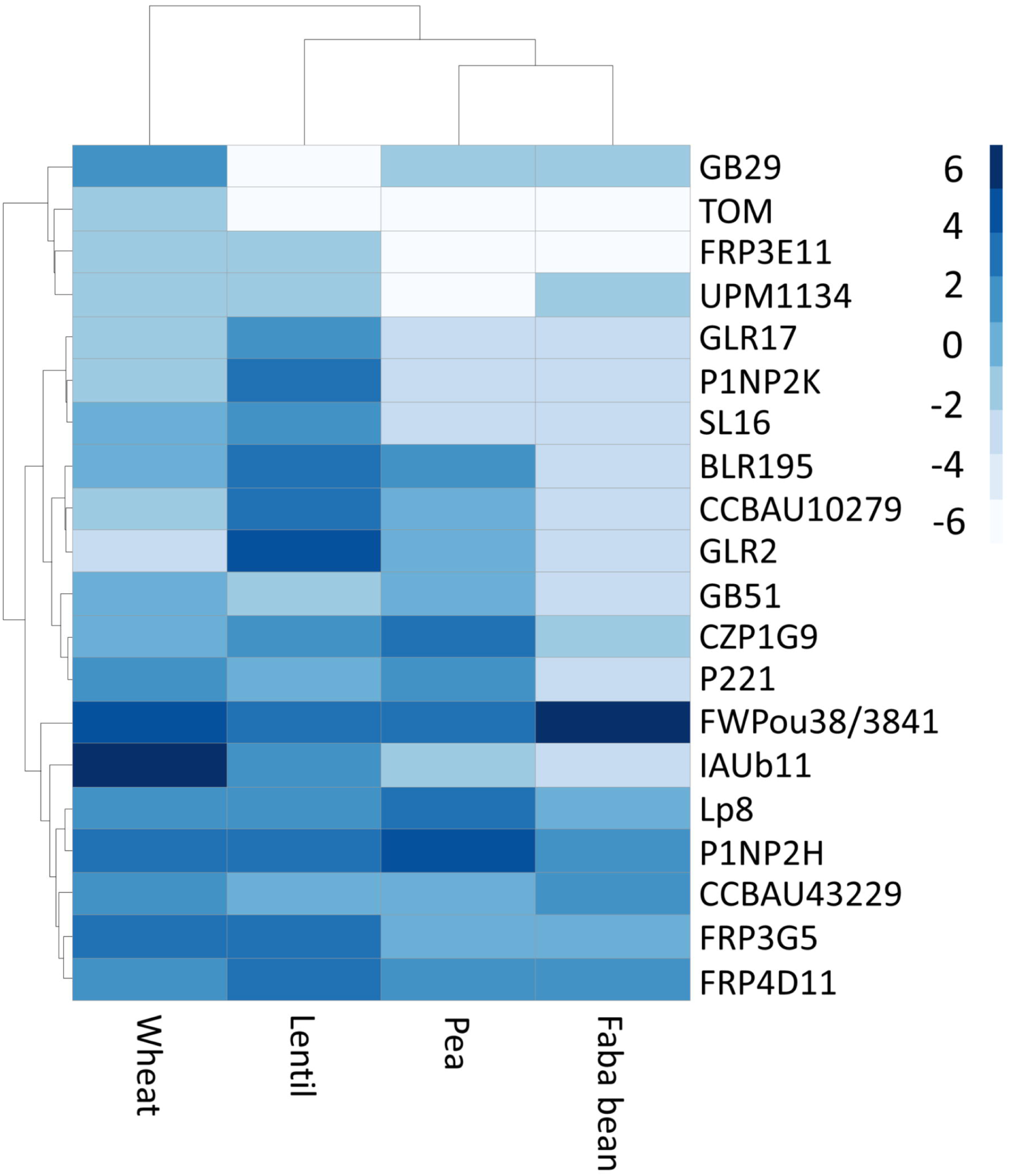
Success of wheat and legume plant colonization in the *Rlv* co-inoculation assays. Hierarchical clustering and heatmap based on the relative abundance values (log2) of each *Rlv* found in the wheat roots and legume nodules during co-inoculation assays (Table S2 and Boivin *et al*., 2020b). FWPou38 or 3841 strains were used in the co-inoculation assays in wheat or legumes respectively as they are undistinguishable by Illumina *nodD* sequencing. Wheat cv. ‘Energo’, Lentil cv. ‘Rosana’, Pea cv. ‘Kayanne’, and Fababean cv. ‘Diva’.

### Different mechanisms could influence *Rlv* competitiveness for endophytic and epiphytic wheat root colonization

The colonization success of a given strain could be due to its ability to colonize the host and/or to compete with other microorganisms. Although we have not tested individually all the *Rlv* strains used in the co-inoculation assays, we have not found any difference for epiphytic growth between the 4 strains we have tested individually. This suggests that the differences observed for the epiphytic growth in the co-inoculation assay are due to differences in competitiveness. Competition for nutrient resources is the most common mechanism underlying variations in bacterial root colonization, as strains/species carrying resource-specific pathways are often those able to rapidly grow and dominate niches (Simons *et al*., 1996; Pieterse *et al*., 2014; Yang *et al*., 2019). Other hypotheses to explain different bacterial abilities to compete for root colonization include antagonism *via* the production of secondary metabolites (Weller *et al*., 2002). However, the minor differences found in this study among the *Rlv* strains in term of metabolic pathways (Fig. S6) did not correlate with the differences observed for wheat root colonization in the co-inoculation assays. Comparative genomics revealed that a cluster of 4 genes annotated as a putative sulfide/taurine transporter was only found in the genomes of the strains FWPou15, FWPou32, FWPou38 and IAUb11 and not in the 21 other strains (gene identifiers in strain IAUb11: RLVIAUB1145.7 - RLVIAUB1145.10). It is possible that this transporter is involved in competitiveness of the FWPou32, FWPou38 and IAUb11 strains by facilitating export of sulfate or sulfite (Weinitschke *et al*., 2007).

Overall, in the co-inoculation assays performed in this study, the competitiveness pattern for epiphytic colonization is similar to the pattern found for endophytic colonization. This indicates that *Rlv* multiplication on the root surface is an important factor underlying the endophytic success of *Rlv* strains. Nevertheless, mechanisms underlying competitiveness of the *Rlv* strains in colonizing the inner part of wheat roots might be different to those regulating the epiphytic growth. First, we found differences for endophytic colonization between the strains we tested individually. Secondly, the endophytic/epiphytic colonization ratio were higher for wheat strains FWPou32 and FWPou38 than for the pea strain IAUb11. The ability of strains to escape recognition by host and to avoid plant defenses might also account for the ability to multiply inside the roots (Pieterse *et al*., 2014). A combination of mechanisms might thus modulate the ability of *Rlv* strains to endophytically colonize wheat roots and explain adaptation of some *Rlv* strains to wheat.

Surprisingly, while almost clonal, FWPou15 and FWPou38 strains showed contrasted ability to colonize wheat roots. Based on short read Illumina genome sequencing, we found only one non-synonymous mutation between FWPou15 and FWPou38 strains, which occurs in a gene encoding a DNA polymerase IV involved in DNA repair. It is known in bacteria, that in response to stress DNA polymerase can synthesize error-containing DNA leading to the formation of genetic variants and different phenotypes of the progeny (Foster, 2005). This process called phenotypic plasticity also leads to bacterial cells showing different responses when inoculated in a plant host (i.e. loss of pathogenicity or decrease of epiphytic growth; Bartoli *et al*., 2015). Further experiments are required to determine whether this mutation, genome re-arrangements and/or epigenetic modifications are responsible for these phenotypic differences. Moreover, these variants represent a promising material to decipher the molecular mechanisms controlling the ability of *Rlv* strains to colonize wheat roots.

### Common molecular mechanisms might be involved in the stimulation of wheat root development and AMF colonization

Several bacteria displaying PGP activity, including rhizobia, have been shown to enhance wheat interaction with AMF (Germida & Walley, 1996; Russo *et al*., 2005; Pivato *et al*., 2009). Soil microorganisms can modulate the auxin-dependent root developmental program leading to lateral root formation in both monocots and dicots (Contreras-Cornejo *et al*., 2009; Pieterse *et al*., 2014). Indeed many PGP bacteria, including rhizobia, produce auxins (Camerini *et al*., 2008; Spaepen & Vanderleyden, 2011; Boivin *et al*., 2016). The effect of *Rlv* on wheat roots development could be attributed to auxin produced by *Rlv* strains. In support of this hypothesis, key enzymes of 3 of the known bacterial auxin synthesis pathways using tryptophan as substrate, the indole acetamide, the indole pyruvate and the tryptamine pathways (Spaepen & Vanderleyden, 2011), are encoded in the FWPou15, FWPou38 and IAUb11 genomes (Fig. S8). Alternatively, rhizobial LCOs can also affect root development both in legumes and non-legumes through the regulation of the plant auxin homeostasis (Herrbach *et al*., 2017; Buendia *et al*., 2019a). Stimulation of AMF colonization by *Rlv* can be indirectly favored by the increase in the number of lateral roots, preferential sites for AMF colonization (Gutjahr *et al*., 2013). LCOs and auxin have been also both shown to stimulate AMF colonization in legumes and non-legumes plants (Maillet *et al*., 2011; Etemadi *et al*., 2014; Buendia *et al*., 2019b) and may also directly affect AM. Effects of exogenous application of either auxin or LCO on root development are dependent on their concentration (Herrbach *et al*., 2017; Buendia *et al*., 2019a). Different levels of auxin and/or LCO production could, at least partially, explain the difference between *Rlv* strains on root development and AMF colonization.

### Levels of endophytic colonization rather than epiphytic colonization might be critical for *Rlv* ability to induce plant responses

Both co-inoculation and single strain inoculation assays showed that FWPou38 and IAUb11 are efficient wheat endophytic colonizers with ∼10^4^ cfu/mg of root tissues on both Energo and Numeric varieties. The bacterial population sizes found in wheat roots for these strains are similar to those previously reported for PGP rhizobia colonizing rice roots (Chaintreuil *et al*., 2000; Mitra *et al*., 2016). Both strains can increase the number of lateral roots and stimulate AMF colonization in Energo. This was not observed on the Numeric variety, suggesting that bacterial effects are plant genotype dependent. The effect of wheat genotype on the colonization by PGP rhizobacteria has been described, for example for bacteria of the genus *Pseudomonas* (Valente *et al*., 2020). Here, we show that beside the colonization level, host responses also vary depending on the plant genotype. Interestingly, FWPou15, had a stimulation activity on Numeric root architecture in which it is an intermediate endophytic colonizer with ∼10^3^ cfu/mg, while not in Energo in which it is a poor endophytic colonizer with ∼10^2^ cfu/mg. Similarly, the poorest endophytic colonizer in both Energo and Numeric, A34, did not induce any root architecture or AMF colonization responses. In this regard, we can hypothesize that *Rlv* strains with the best endophytic colonization ability were able to reach an endophytic population size sufficient for production of auxins, LCOs or other PGP molecules at concentrations required for stimulating lateral root development and AMF colonization. Further analysis with a larger number of strains should establish whether there is a correlation between endophytic colonization ability and stimulation of plant responses and its genotype-genotype dependency.

In conclusion, our study suggest that *Rlv* competitiveness and level of endophytic colonization are critical for potential PGP activities. These novel concepts will help in understanding and/or design microbial consortia based on beneficial bacteria for improving wheat yield/quality in a context of reducing the use of inputs.

## Acknowledgements

We thank Adrien Jallais, Thomas Py and Camille Ribeyre for their support during the wheat phenotypic assays, Dr. Dorian Guetta, who initiated the *in vitro* wheat root colonization studies, and the CREABio Research Station (Auch, France) / Enguerrand Burel and Laurent Escalier who allowed us to sample wheat and provided data about soil composition of the field plots. We also thank Cecile Pouzet from the FR AIB imaging platform for her technical support in analyzing wheat roots colonized with GFP-tagged strains and Julie Cullimore for critical reading of the manuscript. This work was supported by the projects IDEX UNITI “RHIZOWHEAT”, INRAE Department of Plant Health and Environment (SPE) “DIBAM”, ANR ‘‘WHEATSYM’’ (ANR-16-CE20-0025-01), ANR GRaSP (ANR-16-CE20-0021-03) and Laboratoire d’Excellence (LABEX) TULIP (ANR-10-LABX-41).

## Author Contributions

Conceptualization of the project: CB, SB, BL, ML and CMB. Experimental design: CB, SB, BL and ML. Field sampling and microbiology: CG and CB. Wheat phenotyping: CB, VG and MG. Molecular analysis: CB and MG. Statistical Analysis: CB. Microscopy: MM and MCA. Bio-informatic analysis: SB, AC and LC. Manuscript writing: CB, SB, BL, ML and CMB.

## Supporting information

Fig. S1. Soil composition of field plots in which wheat varieties were sampled.

Fig. S2. Images of wheat plantlets grown in gnotobiotic conditions.

Fig. S3. *Rlv* Neighbor-Joining tree based on a portion of the *nodD* gene.

Fig. S4. Nodule sections obtained from vetch plants inoculated with *Rlv* strains isolated from wheat roots.

Fig. S5. Nodulated faba bean root systems inoculated with *Rlv* strains isolated from wheat roots.

Fig. S6. *Rlv* metabolic pathway reconstructions.

Fig. S7. Correlation between the total root length and the lateral root number measured in wheat plantlets inoculated by *Rlv* strains.

Fig. S8. Metabolic reactions of bacterial tryptophan metabolism found in the FWPou15, FWPou38 and IAUb11 strains.

Table S1. List of the wheat varieties sampled and the *Rlv* strains isolated.

Table S2. *gyrB* sequences obtained for strains isolated from wheat roots and the A34 strain.

Table S3. List of the *Rlv* strains used in the co-inoculation assays.

Table S4. Quality information on the co-inoculation assays.

Table S5. Results of the co-inoculation assays.

Table S6. Experimental design for the root colonization in single strain inoculation assays.

Table S7. Experimental design for the root development assays.

Table S8. Experimental design for the mycorrhiza assays.

Table S9. Relative abundances (from linear-mixed model) of the *Rlv* strains in roots during the co-inoculation assays.

Table S10. Numbers of colony-forming units (cfu) in the single strain inoculation assays.

Table S11. Total root lengths and lateral root numbers in the root development assays.

Table S12. Number of colonization sites in the mycorrhiza assays.

Table S13. Statistical analysis of the co-inoculation assays.

Table S14. Statistical analysis of the in the single strain inoculation assays.

Table S15. Statistical analysis of the root development assays.

Table S16. Statistical analysis of the mycorrhiza assays. Supplementary methods (S1-S9).

## References

Arkin AP, R., Henry CS, Harris NL, Stevens RL, Maslov S, Dehal P, Ware D, Perez F et al. 2018. KBase: the united states department of energy systems biology knowledgebase. Nature biotechnology 36: 566–569.

Barker D, Pfaff T, Moreau D, Groves E, Ruffel R, Lepetit M, Whiteh S, Maillet F, Ramakrishnan MN, Journet EP. 2006. Growing *M. truncatula*: choice of substrates and growth conditions. In: Medicago truncatula handbook. (eds U. Mathesius, E. Journet, & L. Sumner), pp. http://www.noble.org/MedicagoHandbook/. ISBN 0-9754303-1-9.

Bartoli C, Lamichhane JR, Berge O, Varvaro L, Morris CE. 2015. Mutability in *Pseudomonas viridiflava* as a programmed balance between antibiotic resistance and pathogenicity. Molecular Plant Pathology 16: 860–869.

Beringer JE. 1974. R factor transfer in *Rhizobium leguminosarum*. Microbiology 84: 188–198.

Biswas JC, Ladha JK, Dazzo FB, Yanni YG, Rolfe BG. 2000. Rhizobial inoculation influences seedling vigor and yield of rice. Agronomy Journal 92: 880–886.

Boivin S, Fonouni-Farde C, Frugier F. 2016. How auxin and cytokinin phytohormones modulate root microbe interactions. Frontiers in Plant Science 7: 1240.

Boivin S, Ait Lahmidi N, Sherlock D, Bonhomme M, Dijon D, Heulin-Gotty K, Le-Quer, A, Pervent M, Tauzin M et al. 2020a. Host - specific competitiveness to form nodules in *Rhizobium leguminosarum* symbiovar *viciae*. New Phytologist 266: 555–568

Boivin S, Mahé F, Pervent M, Tancelin M, Tauzin M, Wielbo J, Mazurier S, Young PJ, Lepetit M. 2020b. Genetic variation in host-specific competitiveness of the symbiont *Rhizobium leguminosarum* symbiovar *viciae*. Authorea. doi: 10.22541/au.159237007.72934061.

Bonfante P, Genre A. 2010. Mechanisms underlying beneficial plant–fungus interactions in mycorrhizal symbiosis. Nature Communications 1: 48.

Brettin T, Davis JJ, Disz T, Edwards RA, Gerdes S, Olsen GJ, Olson R, Overbeek R, Parrello B, Pusch GD et al. 2015. RASTtk: A modular and extensible implementation of the RAST algorithm for building custom annotation pipelines and annotating batches of genomes. Scientific Reports 5: 8365.

Buendia L, Maillet F, O’Connor D, van de-Kerkhove Q, Danoun S, Gough C, Lefebvre B, Bensmihen S. 2019a. Lipo-chitooligosaccharides promote lateral root formation and modify auxin homeostasis in *Brachypodium distachyon*. New Phytologist 221: 2190–2202.

Buendia L, Ribeyre C, Bensmihen S, Lefebvre B. 2019b. *Brachypodium distachyon* tar2lhypo mutant shows reduced root developmental response to symbiotic signal but increased arbuscular mycorrhiza. Plant Signaling and Behavior 14: e1651608.

Camerini S, Senatore B, Lonardo E, Imperlini E, Bianco C, Moschetti G, Rotino GL, Campion B, Defez R. 2008. Introduction of a novel pathway for IAA biosynthesis to rhizobia alters vetch root nodule development. Archives of Microbiology 190: 67–77.

Catoira R, Galera C, de Billy F, Penmetsa RV, Journet E-P, Maillet F, Rosenberg C, Cook D, Gough C, Dénarié J. 2000. Four genes of Medicago truncatula controlling components of a Nod factor transduction pathway. Plant Cell 12: 1647–1666.

Chaintreuil C, Giraud E, Prin Y, Lorquin J, Bâ A, Gillis M, de Lajudie P, Dreyfus B. 2000. Photosynthetic bradyrhizobia are natural endophytes of the african wild rice *Oryza breviligulata*. Applied and Environmental Microbiology 66: 5437–5447.

Contreras-Cornejo HA, Macías-Rodríguez L, Cortés-Penagos C, López-Bucio J. 2009. *Trichoderma virens*, a plant beneficial fungus, enhances biomass production and promotes lateral root growth through an auxin-dependent mechanism in *Arabidopsis*. Plant Physiology 149: 1579–1592.

Depret G, Houot S, Allard MR, Breuil MC, Nouaïm R, Laguerre G. 2004. Long-term effects of crop management on *Rhizobium leguminosarum* biovar *viciae* populations. FEMS Microbiology Ecology 51: 87–97.

Dray S, Dufour A. 2007. The ade4 package: implementing the duality diagram for ecologists. Journal of Statistical Software 22: 1–20.

Emms DM, Kelly S. 2019. OrthoFinder: phylogenetic orthology inference for comparative genomics. Genome Biology 20: 238.

Etemadi M, Gutjahr C, Couzigou JM, Zouine M, Lauressergues D, Timmers A, Audran C, Bouzayen M, Bécard G, Combier JP. 2014. Auxin perception is required for arbuscule development in arbuscular mycorrhizal symbiosis. Plant Physiology 166: 281–292.

Foster PL. 2005. Stress responses and genetic variation in bacteria. Mutation Research 569: 3–11.

Germida JJ, Walley FL. 1996. Plant growth-promoting rhizobacteria alter rooting patterns and arbuscular mycorrhizal fungi colonization of field-grown spring wheat. Biology and Fertility of Soils. 23: 113–120.

Girardin A, Wang T, Ding Y, Keller J, Buendia L, Gaston M, Ribeyre C, Gasciolli V, Auriac MC, Vernié T, et al. 2019. LCO receptors involved in arbuscular mycorrhiza are functional for rhizobia perception in legumes. Current Biology 29: 4249-4259.e5.

Götz R, Evans IJ, Downie JA, Johnston AWB. 1985. Identification of the host-range DNA which allows *Rhizobium leguminosarum* strain TOM to nodulate cv. Afghanistan peas. Molecular and General Genetics 201: 296–300.

Gutjahr C, Paszkowski U. 2013. Multiple control levels of root system remodeling in arbuscular mycorrhizal symbiosis. Frontiers in Plant Science 4: 204.

Haichar F el Z, Santaella C, Heulin T, Achouak W. 2014. Root exudates mediated interactions belowground. Soil Biology and Biochemistry 77: 69–80.

Henry CS, DeJongh M, Best AA, Frybarger PM, Linsay B, Stevens RL. 2010. High-throughput generation, optimization and analysis of genome-scale metabolic models. Nature Biotechnology 28: 977–982.

Herrbach V, Chirinos X, Rengel D, Agbevenou K, Vincent R, Pateyron S, Huguet S, Balzergue S, Pasha A, Provart N, et al. 2017. Nod factors potentiate auxin signaling for transcriptional regulation and lateral root formation in *Medicago truncatula*. Journal of Experimental Botany 68: 569–583.

Hilali A, Prevost D, Broughton WJ, Antoun H. 2001. Effects of inoculation with *Rhizobium leguminosarum* biovar *trifolii* on wheat cultivated in clover crop rotation agricultural soil in Morocco. Canadian Journal of Microbiology 47: 590–593.

Höflich G, Wiehe W, Hecht-Buchholz C. 1995. Rhizosphere colonization of different crops with growth promoting *Pseudomonas* and *Rhizobium* bacteria. Microbiological Research 150: 139–147.

Hogland DR, Arnon DI. 1950. The water culture method for growing plants without soil. 1950. University of California, College of Agriculture, Agricultural Experiment Station. Berkeley, California. Vol.347 No.2nd edit pp.32 pp.

Hucka M, Finney A, Sauro HM, Bolouri H, Doyle JC, Kitano H, Arkin AP, Bornstein BJ, Bray D, Cornish-Bowden A, et al. 2003. The systems biology markup language (SBML): a medium for representation and exchange of biochemical network models. Bioinformatics 19: 524–531.

Huerta-Cepas J, Forslund K, Coelho LP, Szklarczyk D, Jensen LJ, von Mering C, Bork P. 2017. Fast genome-wide functional annotation through orthology assignment by eggNOG-Mapper. Molecular Biology and Evolution 34: 2115–2122.

Machado D, Andrejev S, Tramontano M, Patil KR. 2018. Fast automated reconstruction of genome-scale metabolic models for microbial species and communities. Nucleic Acids Research 46: 7542–7553.

Maillet F, Poinsot V, André O, Puech-Pagès V, Haouy A, Gueunier M, Cromer L, Giraudet D, Formey D, Niebel A, et al. 2011. Fungal lipochitooligosaccharide symbiotic signals in arbuscular mycorrhiza. Nature 469: 58–63.

Martens M, Dawyndt P, Coopman R, Gillis M, De Vos P, Willems A. 2008. Advantages of multilocus sequence analysis for taxonomic studies: a case study using 10 housekeeping genes in the genus *Ensifer* (including former *Sinorhizobium*). International Journal of Systamaic and Evolutionary Microbiology 58: 200–214.

Mitra S, Mukherjee A, Wiley-Kalil A, Das S, Owen H, Reddy PM, Ané J-M, James EK, Gyaneshwar P. 2016. A rhamnose-deficient lipopolysaccharide mutant of Rhizobium Sp. IRBG74 is defective in root colonization and beneficial interactions with its flooding-tolerant hosts *Sesbania cannabina* and wetland rice. Journal of Experimental Botany 67: 5869–5884.

Oono R, Denison RF. 2010. Comparing symbiotic efficiency between swollen versus nonswollen rhizobial bacteroids. Plant Physiology 154: 1541–1548.

Ortíz-Castro R, Contreras-Cornejo HA, Macías-Rodríguez L, López-Bucio J. 2009. The role of microbial signals in plant growth and development. Plant Signaling and Behavior 4: 701–712.

Peng S, Biswas JC, Ladha JK, Gyaneshwar P, Chen Y. 2002. Influence of rhizobial inoculation on photosynthesis and grain yield of rice. Agronomy Journal 94:925–929.

Pieterse CMJ, Zamioudis C, Berendsen RL, Weller DM, Van Wees SCM, Bakker PAHM. 2014. Induced systemic resistance by beneficial microbes. Annual Review of Phytopathology 52: 347–375.

Pivato B, Offre P, Marchelli S, Barbonaglia B, Mougel C, Lamanceau P, Berta G. 2009. Bacterial effects on arbuscular mycorrhizal fungi and mycorrhiza development as influenced by the bacteria, fungi, and host plant. Mychorriza 19: 81–90.

Pouler S. 1995. Accuracy of measurements with Mac/WinRhizo. Reagent Instrumentstechnical note N°3.

Raklami A, Bechtaoui N, Tahiri AI, Anli M, Meddich A, Oufdou K. 2019. Use of rhizobacteria and mycorrhizae consortium in the open field as a strategy for improving crop nutrition, productivity and soil fertility. Frontiers in Microbiology 10: 1106.

Russo A, Felici C, Toffanin A, Götz M, Collados C, Barea JM, Moënne-Loccoz Y, Smalla K, Vanderleyden J, Nuti M. 2005. Effect of *Azospirillum* inoculants on arbuscular mycorrhiza establishment in wheat and maize plants. Biology and Fertility of Soils 41: 301–309.

Santoyo G, Moreno-Hagelsieb G, del Carmen Orozco-Mosqueda M, Glick BR. 2016. Plant growth-promoting bacterial endophytes. Microbiological Research 183: 92–99.

Simons M, Van Der Bij AJ, Brand I, De Weger LA, Wijffelman CA, Lugtenberg BJJ. 1996. Gnotobiotic system for studying rhizosphere colonization by plant growth-promoting *Pseudomonas* bacteria. Molecular Plant-Microbe Interactions 9: 600–607.

Spaepen S, Vanderleyden J. 2011. Auxin and plant-microbe interactions. Cold Spring Harbor Perspectives in Biology 3: a001438.

Tamura K, Peterson D, Peterson N, Stecher G, Nei M, Kumar S. 2011. MEGA5: Molecular evolutionary genetics analysis using maximum likelihood, evolutionary distance, and maximum parsimony methods. Molecular Biology and Evolution 28: 2731–2739.

Valente J, Gerin F, Le Gouis J, Moënne-Loccoz Y, Prigent–Combaret C. 2020. Ancient wheat varieties have a higher ability to interact with plant growth-promoting rhizobacteria. Plant Cell and Environment 43: 246–260.

Webster G, Gough C, Vasse J, Batchelor CA, O’Callaghan KJ, Kothari SL, Davey MR, Dénarié J, Cocking EC. 1997. Interactions of rhizobia with rice and wheat. Plant and Soil 194: 115–122.

Weinitschke S, Denger K, Cook AM, Smits THM. 2007. The DUF81 protein TauE in Cupriavidus necator H16, a sulfite exporter in the metabolism of C2 sulfonates. Microbiology 153: 3055–3060.

Weller DM, Raaijmakers JM, Gardener BBM, Thomashow LS. 2002. Microbial populations responsible for specific soil suppressiveness to plant pathogens. Annual Review of Phytopathology 40: 309–348.

Xia X. 2017. DAMBE6: New tools for microbial genomics, phylogenetics, and molecular evolution. Journal of Heredity 108: 431–437.

Yang C, Dong Y, Friman VP, Jousset A, Wei Z, Xu Y, Shen Q. 2019. Carbon resource richness shapes bacterial competitive interactions by alleviating growth-antibiosis trade-off. Functional Ecology 33: 868–875.

Yanni YG, Dazzo FB,. 2010. Enhancement of rice production using endophytic strains of *Rhizobium leguminosarum* bv. *trifolii* in extensive field inoculation trials within the Egypt Nile delta. Plant and Soil 366: 129–141.

Zeze A, Mutch LA, Young PW. 2001. Direct amplification of *nodD* from community DNA reveals the genetic diversity of *Rhizobium leguminosarum* in soil. Environmental Microbiology 3: 363–370.

