## Supplementary Information for "*Rhizobium leguminosarum* symbiovar *viciae* strains are natural wheat endophytes and can stimulate root development and colonization by arbuscular mycorrhizal fungi"

***New Phytologist* Supporting Information**

The following Supporting Information is available for this article:

### Supplemental Figures

**Fig. S1. Soil composition of field plots in which wheat varieties were sampled.** Values are expressed as percentages of each soil component found in 2 plots LH7 (on the left panel) and LH1 (on the right panel). See Table S1 for varieties grown in each plot. Soil analyses were performed on different samples obtained at a depth ranging from 0 to 40 cm. Values for each depth range (0-15 and 15-40 cm) were averaged to calculate the percentage of each soil component.

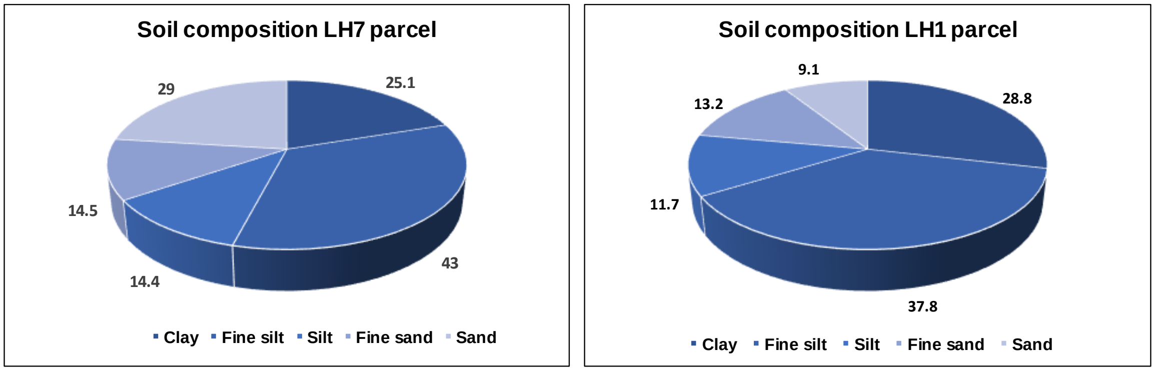


**Fig. S2. Images of wheat plantlets grown in gnotobiotic conditions.** (a) 110 ml tubes and (b) 20×20 cm plates.

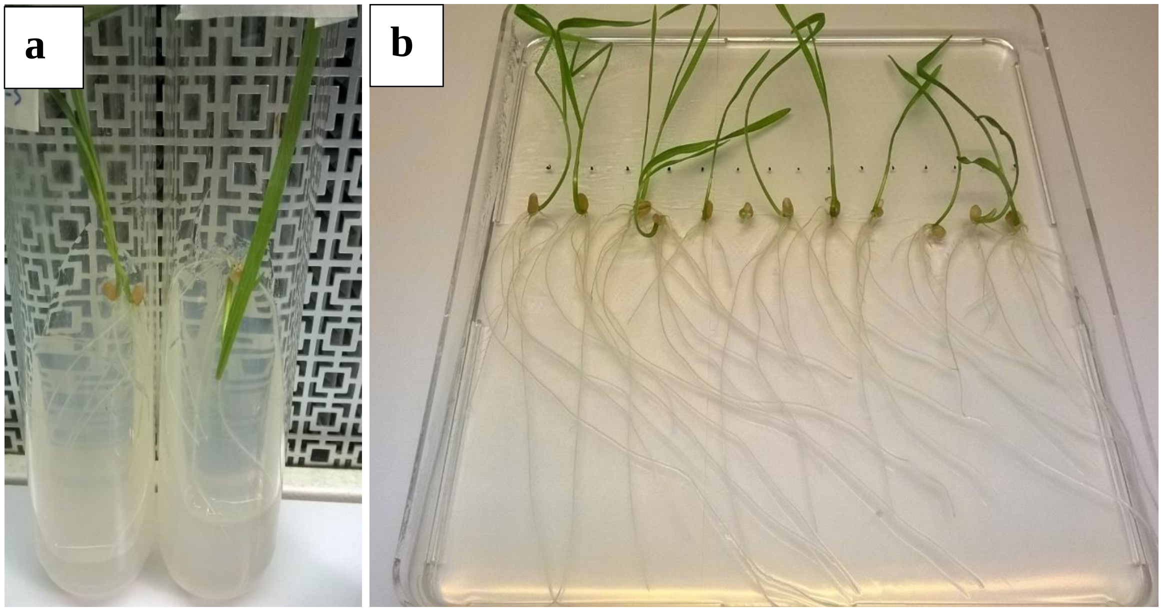

**Fig. S3. *Rlv* Neighbor-Joining tree based on a portion of the *nodD* gene (605 bp).** It includes 3 *Rlv* strains isolated from wheat roots (indicated in blue), *Rlv* strains representative of the Nod groups (Boivin *et al*., 2020) and the control strain A34 (indicated in green). WSM1689 (*R. leguminosarum* sv. *trifolii*) and 1021 (*E. meliloti*) have been used as outgroups.

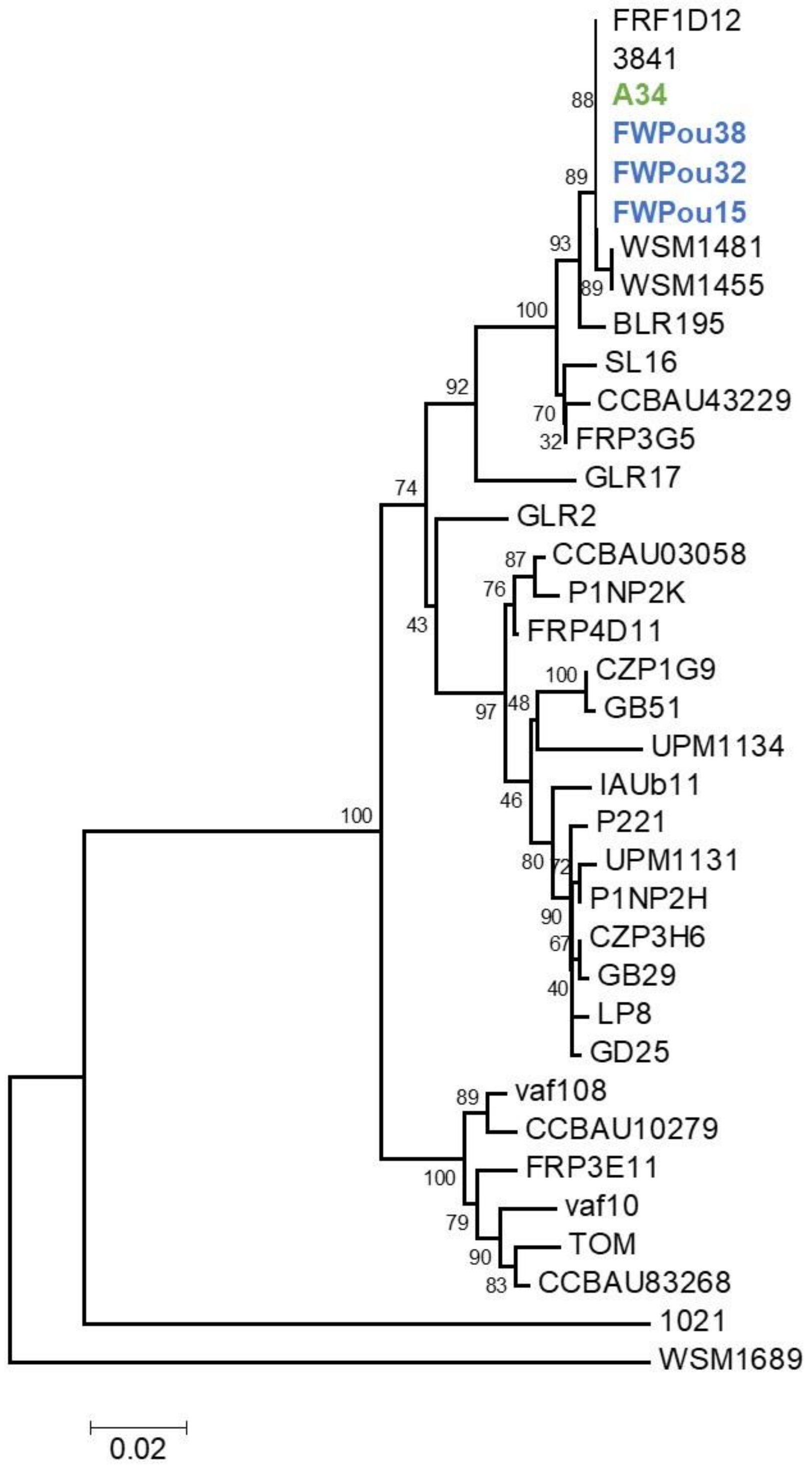

**Fig. S4. Nodule sections obtained from vetch plants inoculated with *Rlv* strains isolated from wheat roots**. (a) FWPou15 (b) FWPou32, (c) FWPou38 and (d) A34 (control). Root systems were imaged 6 weeks post-inoculation. Scale bars represent 100µm.

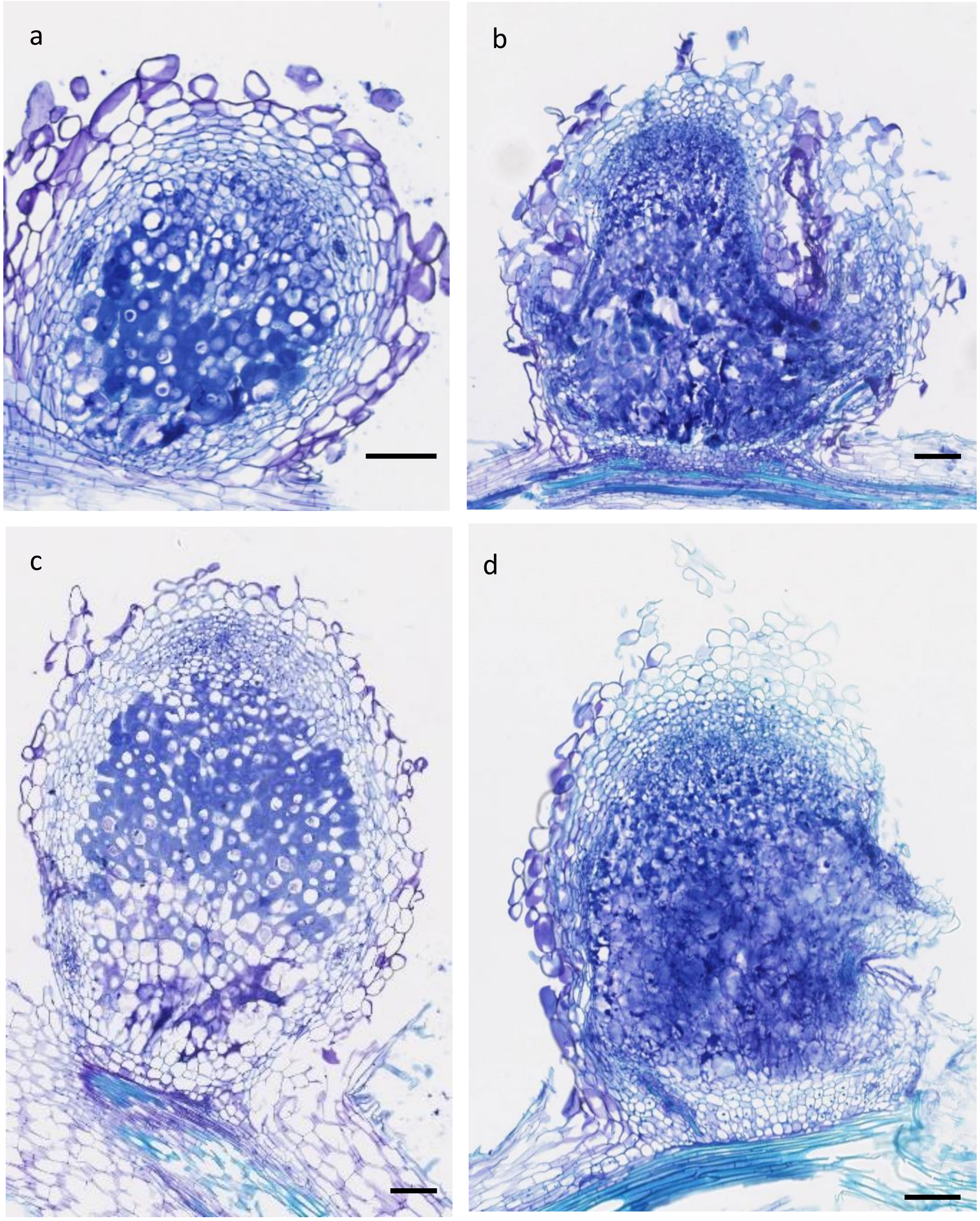

**Fig. S5. Nodulated faba bean root systems inoculated with *Rlv* strains isolated from wheat roots**. (a) FWPou15 (b) FWPou32, (c) FWPou38 and (d) A34 (control). Root systems were imaged 6 weeks post-inoculation.

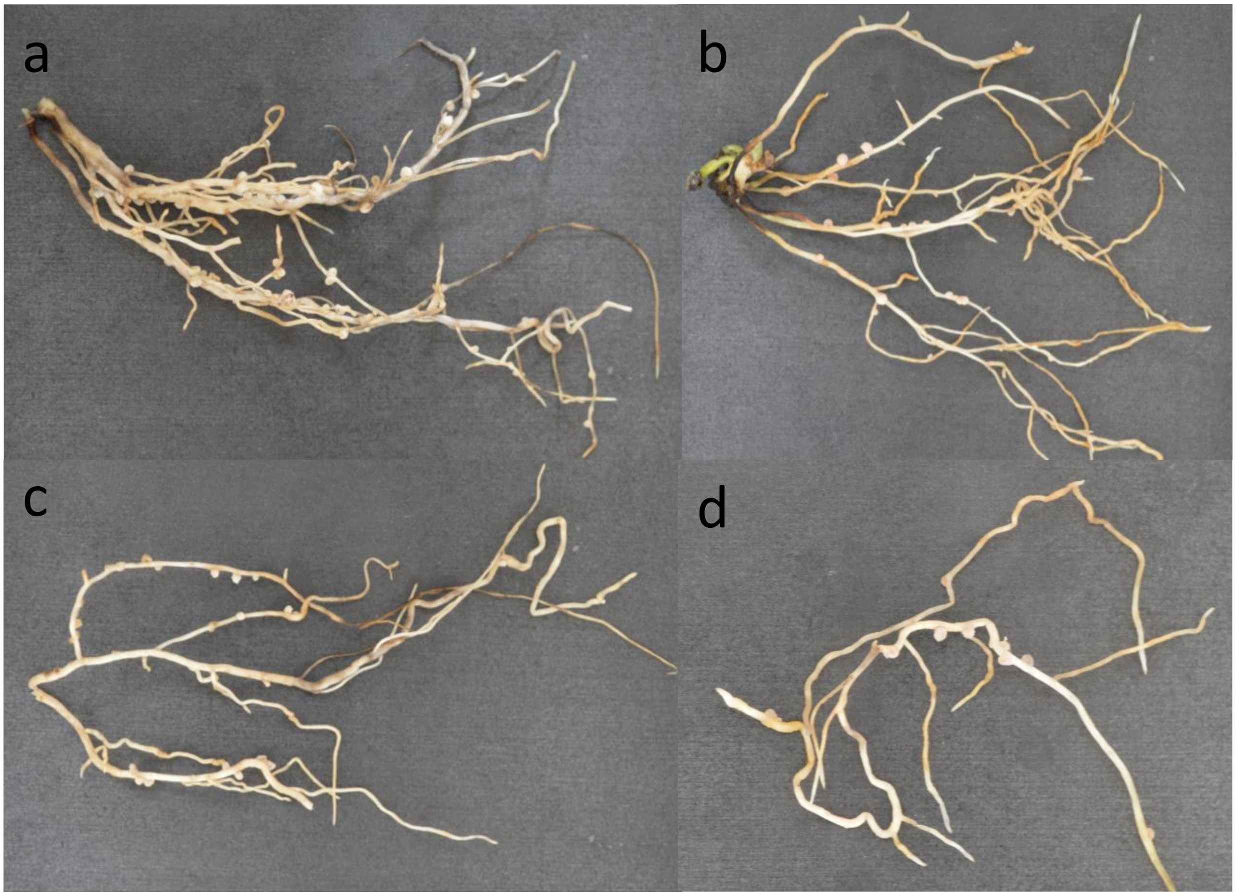

**Fig. S6. *Rlv* metabolic pathway reconstructions.**

(a) Intersection graphic representing the number of metabolic reactions shared among the *Rlv* strains isolated from wheat and strains representing the genetic diversity of the *Rlv* species complex (Table S2). The number of reactions indicated on the top *x*-axis (among a total of 2260 identified reactions) are shared between the strains indicated by black dots in the vertical lines. All strains share 1967 reactions (first vertical line on the left). Other reactions are shared by subsets of strains. Set size (on the left) indicates the number of identified reactions in each strain. (b) Correspondence Analysis (CA) inferred on the metabolic reactions identified in the same *Rlv* strains as in (a).

a

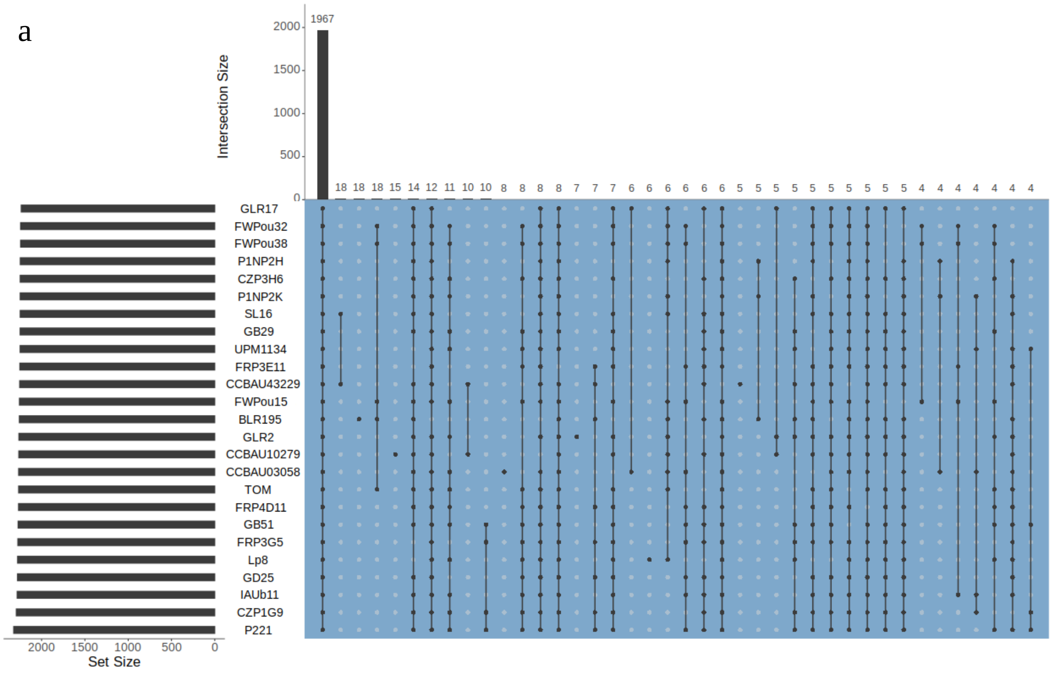


b

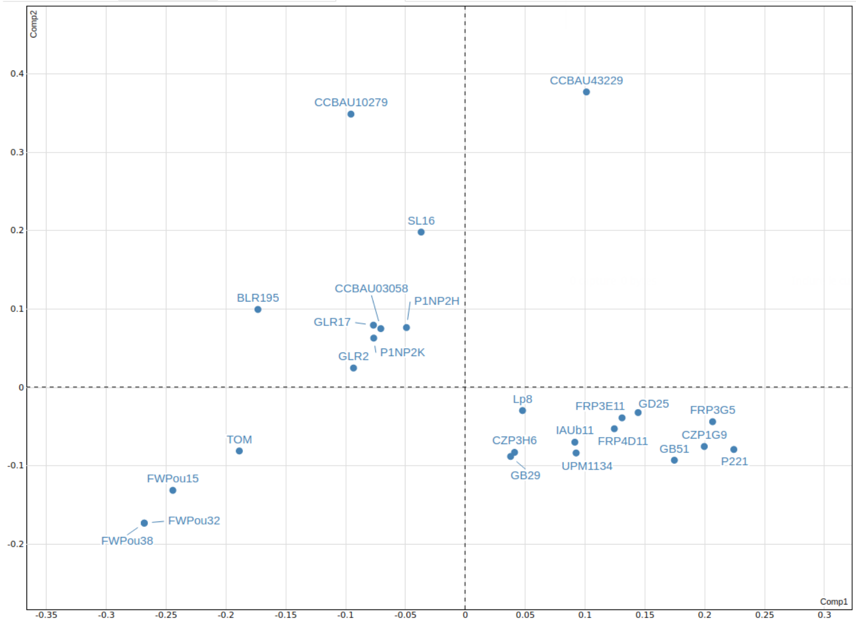

**Fig. S7. Correlation between the total root length and the lateral root number measured in wheat plantlets inoculated by *Rlv* strains**. Data obtained on the wheat varieties Energo (a) and Numeric (b) 12 dpi with the strains A34, FWPou15, IAUb11 and FWPou38 (pooled in the plot). *x*-axes represent the total root length and *y*-axes represent the number of later roots obtained in the same experiments. $R^2$ showed in the graphs indicates a high correlation between the 2 factors.

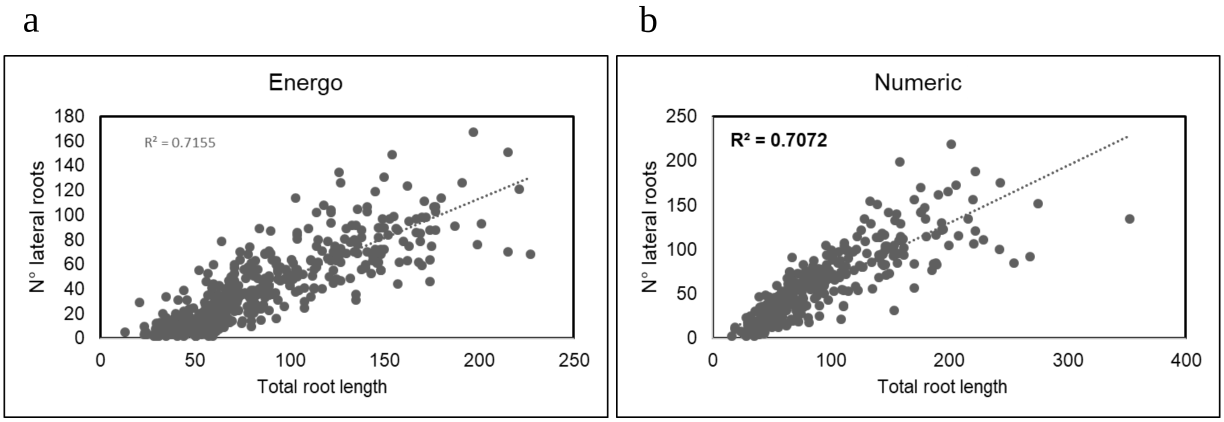

**Fig. S8. Metabolic reactions of bacterial tryptophan metabolism found in the FWPou15, FWPou38 and IAUb11 strains**. Genes encoding the enzymes highlighted in blue are found in the FWPou15 and FWPou38 (a) and IAUb11 (b) genomes. Indole acetic acid (IAA) is colored in red. Enzymes involved in the indole acetamide (green), indole pyruvate (blue) and tryptamine (brown) pathways leading to auxin IAA are shown. The tryptophan metabolism was reconstructed using kbase.

a

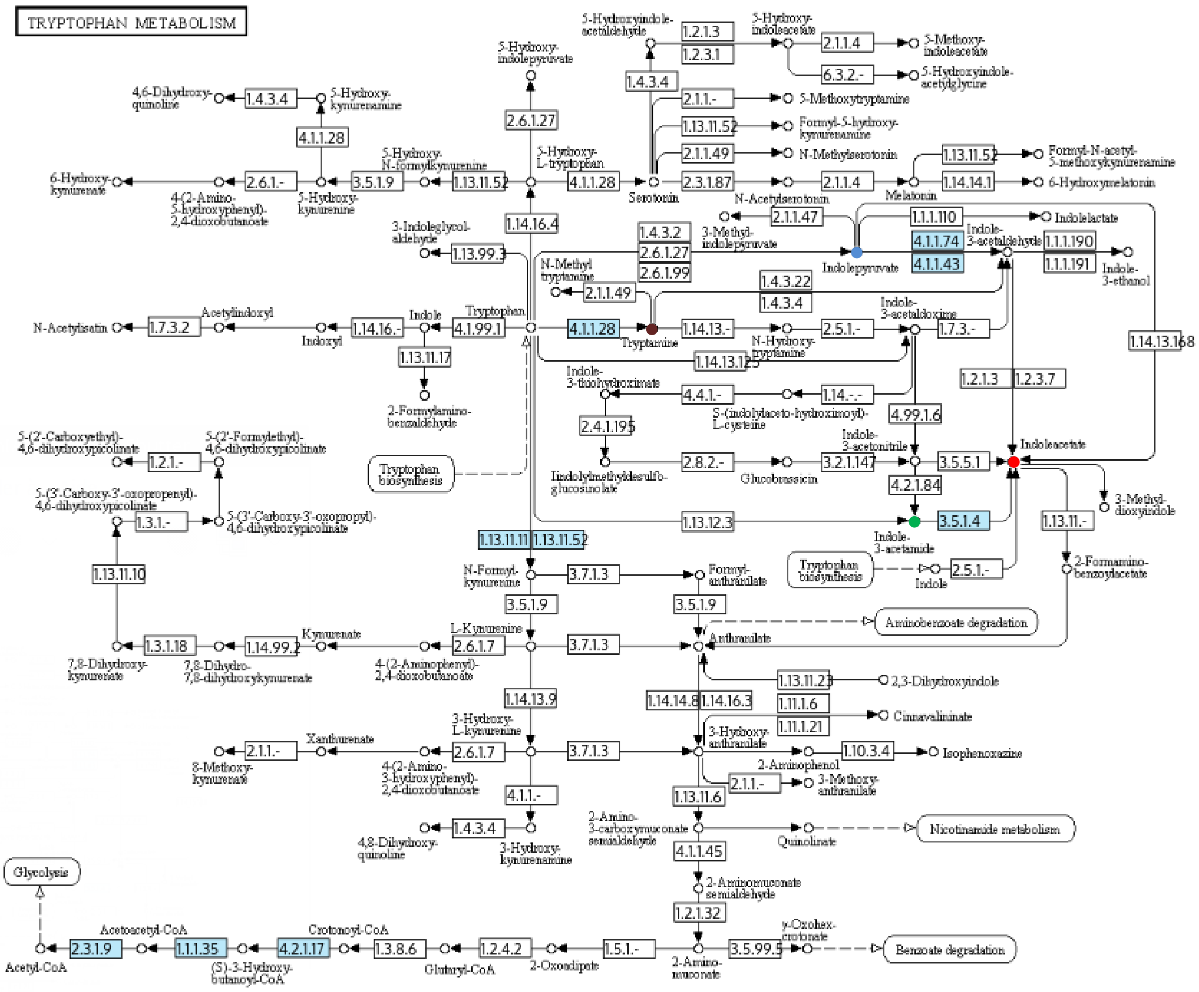

b

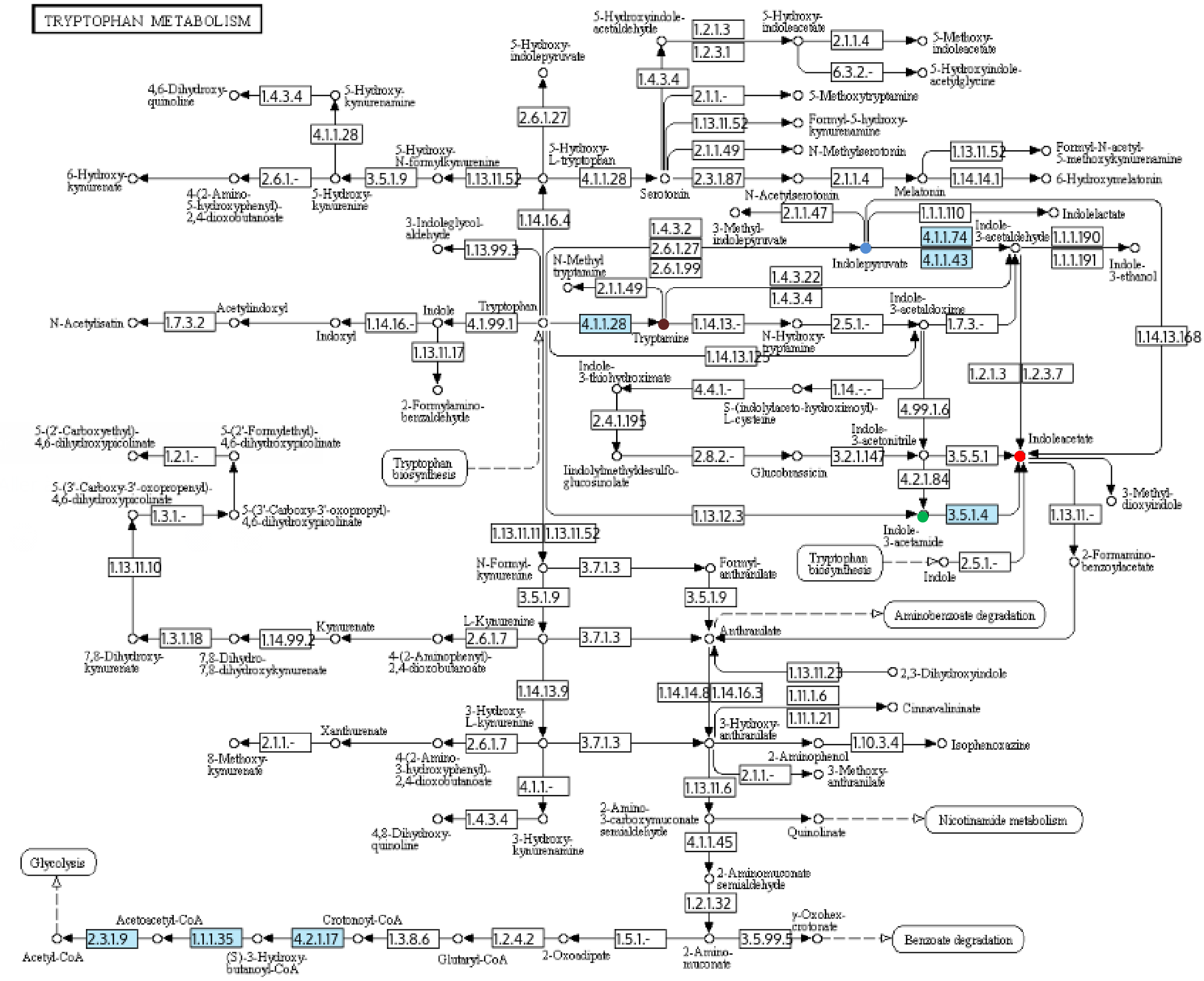
TRYPTOPHAN METABOLISM
5-Hydroxyindole-acetaldehyde
1.2.1.3
1.2.3.1
5-Hydroxy-indoleacetate
2.1.1.4
5-Methoxy-indoleacetate
6.3.2.-
5-Hydroxyindole-acetylglycine
5-Hydroxy-indolepyruvate
1.4.3.4
2.1.1.-
5-Methoxytryptamine
1.13.11.52
Formyl-5-hydroxy-kynurenamine
2.1.1.49
N-Methylserotonin
4,6-Dihydroxy-quinoline
1.4.3.4
5-Hydroxy-kynurenamine
4.1.1.28
2.6.1.27
1.13.11.52
Formyl-N-acetyl-5-methoxykynurenamine
6-Hydroxy-kynurenate
2.6.1.-
3.5.1.9
1.13.11.52
4.1.1.28
2.3.1.87
2.1.1.4
1.14.14.1
6-Hydroxymelatonin
4-(2-Amino-5-hydroxyphenyl)-2,4-dioxobutanoate
5-Hydroxy-kynurenine
5-Hydroxy-N-formylkynurenine
5-Hydroxy-L-tryptophan
Serotonin
N-Acetylserotonin
Melatonin
1.14.16.4
2.1.1.47
1.1.1.110
Indolelactate
3-Indoleglycol-aldehyde
1.13.99.3
1.4.3.2
2.6.1.27
2.6.1.99
3-Methyl-indolepyruvate
Indolepyruvate
4.1.1.74
4.1.1.43
Indole-3-acetaldehyde
1.1.1.190
1.1.1.191
Indole-3-ethanol
N-Methyl tryptamine
2.1.1.49
1.4.3.22
1.4.3.4
N-Acetylisatin
1.7.3.2
Acetylindoxyl
Indoxyl
1.14.16.-
Indole
4.1.99.1
Tryptophan
4.1.1.28
Tryptamine
1.14.13.-
N-Hydroxy-tryptamine
2.5.1.-
Indole-3-acetaldoxime
1.7.3.-
1.14.13.168
1.13.11.17
1.14.13.125
2-Formylamino-benzaldehyde
Indole-3-thiohydroximate
4.4.1.-
S-(indolylaceto-hydroximoyl)-L-cysteine
1.14.-.-
1.2.1.3
1.2.3.7
2.4.1.195
4.99.1.6
5-(2'-Carboxyethyl)-4,6-dihydroxypicolinate
1.2.1.-
5-(2'-Formylethyl)-4,6-dihydroxypicolinate
Tryptophan biosynthesis
Indolylmethylthiohydroximate glucosinolate
2.8.2.-
Glucobrassicin
3.2.1.147
Indole-3-acetonitrile
3.5.5.1
Indoleacetate
5-(3'-Carboxy-3'-oxopropenyl)-4,6-dihydroxypicolinate
1.3.1.-
5-(3'-Carboxy-3'-oxopropyl)-4,6-dihydroxypicolinate
4.2.1.84
3-Methyl-dioxyindole
1.13.12.3
3.5.1.4
Indole-3-acetamide
1.13.11.-
1.13.11.11
1.13.11.52
Tryptophan biosynthesis
Indole
2.5.1.-
2-Formamino-benzoylacetate
1.13.11.10
N-Formyl-kynurenine
3.7.1.3
Formyl-anthranilate
3.5.1.9
3.5.1.9
Aminobenzoate degradation
Kynurenate
L-Kynurenine
2.6.1.7
3.7.1.3
Anthranilate
7,8-Dihydroxy-kynurenate
1.3.1.18
7,8-Dihydro-7,8-dihydroxykynurenate
1.14.99.2
4-(2-Aminophenyl)-2,4-dioxobutanoate
1.13.11.23
2,3-Dihydroxyindole
1.14.13.9
1.14.14.8
1.14.16.3
1.11.1.6
1.11.1.21
Cinnavalininate
Xanthurenate
3-Hydroxy-L-kynurenine
3-Hydroxy-anthranilate
2.1.1.-
2.6.1.7
3.7.1.3
1.10.3.4
Isophenoxazine
2-Aminophenol
8-Methoxy-kynurenate
4-(2-Amino-3-hydroxyphenyl)-2,4-dioxobutanoate
2.1.1.-
3-Methoxy-anthranilate
4.1.1.-
1.13.11.6
1.4.3.4
4,8-Dihydroxy-quinoline
3-Hydroxy-kynurenamine
2-Amino-3-carboxymuconate semialdehyde
Quinolinate
Nicotinamide metabolism
4.1.1.45
Glycolysis
2-Aminomuconate semialdehyde
1.2.1.32
Acetoacetyl-CoA
Crotonoyl-CoA
y-Oxohex-crotonate
Benzoate degradation
2.3.1.9
1.1.1.35
4.2.1.17
1.3.8.6
1.2.4.2
1.5.1.-
3.5.99.5
Acetyl-CoA
(S)-3-Hydroxy-butanoyl-CoA
Glutaryl-CoA
2-Oxoadipate
2-Amino-muconate

**Supplemental methods**

**Methods S1** *Seed sterilization and germination for all plant species used in the study.*

**Sterilization and germination of pea seeds:**

Seeds were incubated for 5 min in ethanol 70%, rinsed 3 times with sterilized water, further incubated for 7 min in a 2.5 % sodium hypochlorite solution and rinsed 3 times in sterilized water. Seeds were then imbibed 2 h in sterilized water and germinated 3 days at 20°C on 4g/L agar plates.

**Sterilization and germination of faba bean and common vetch seeds:**

Seeds were incubated for 20 min in concentrated $H_2SO_4$. After removing the acid, seeds were rinsed 3 times in sterilized water, further incubated for 20 min in a 3.9 % sodium hypochlorite solution, rinsed 3 times in sterilized water. Seeds were then imbibed for 4 h in sterilized water, incubated 4 days at 4°C on 4g/L agar plates and germinated 24 h at 20°C.

**Sterilization and germination of clover seeds:**

Seeds were incubated for 2 min in ethanol 70%, rinsed 3 times with sterilized water, further incubated for 5 min in a 2.5 % sodium hypochlorite solution and rinsed 3 times in sterilized water. Seeds were then germinated 3 days at 20°C on 4g/L agar plates.

**Sterilization and germination of wheat seeds:**

Seeds were incubated for 5 min in ethanol 70 %, rinsed 3 times with sterilized water, further incubated for 45 min in a 2.3 % sodium hypochlorite solution (with constant stirring), and washed five times in sterilized water. Seeds were then incubated 1 week at 4°C on 4g/L agar plates and germinated 2 days at 28 °C.

**Methods S2** *Genome sequencing and bioinformatics analysis.*

To compare the *Rlv* strains isolated from wheat to the diversity of the *Rlv* species complex, bacterial genomes were sequenced by MicrobesNG (Birmingham, UK, https://microbesng.uk/) on an Illumina HiSeq platform using a 2×250bp paired end protocol. Genomic DNA libraries were prepared using Nextera XT Library Prep Kit (Illumina, San Diego, USA). High-quality paired reads were assembled by the Galaxy/BBRIC pipeline (https://bbric-pipelines.toulouse.inra.fr/) and genome annotations were performed using EuGene-PP (Sallet *et* *al.*, 2014). The pairwise ANI values were calculated using the JSpecies software (http://jspecies.ribohost.com/jspeciesws).

Heatmaps were built using the pheatmap R package (Kolde & Vilo, 2015). The nucleotide sequences of the nodABCDEFIJLMN genes were concatenated and aligned using ClustalOmega (https://www.ebi.ac.uk/services), and a Neighbor-Joining (NJ) phylogenetic tree was built using MEGA v7.0.26 (www.megasoftware.net/).

**Methods S3** *Microscopy of nodules and wheat roots inoculated with GFP transformed strains.*

To verify the colonization of nodules by the *Rlv* wheat strains, 3 nodules from pea and faba bean were collected with a sterilized scalpel and fixed with glutaraldehyde 1.25% in 0.1 M PBS pH7.2 for 30 min under vacuum. Fixed nodules were rinsed with sterilized water and dried. Nodules were sectioned after inclusion in 6% agarose low gelling temperature using a vibratome VT 1000S and sections were stained with 5% toluidine blue solution. Stained sections were then analyzed with the NanoZoomer HT (HAMAMATSU).

To investigate the colonization pattern of wheat roots by the FWPou15 GFP-tagged strain, wheat roots were grown as described in the “root colonisation in single strain inoculation assays” section. Roots were excised 7dpi, fixed in paraformaldehyde (PFA) at 3.7% in PBS under vacuum (30 min), washed and analyzed under a confocal scanning laser microscopy (SP8; Leica).

For clearing method, roots were fixed in 4% PFA in PBS solution and incubated in RapiClearR 1.47 for 1 month for clearing. Samples were analyzed with a confocal laser scanning microscope (SP8; Leica) using a ×25 water immersion objective lens (numerical aperture 0.95; PL APO). GFP fluorescence of bacteria was excited at 488 nm and recorded in one of the confocal channels in the 505 to 545 nm emission range. Auto-fluorescence of roots was excited at 405nm and 561 nm and detected respectively in the 415-450nm and 595-620 nm range, respectively.

**Methods S4.** *Production of germ-free wheat plants for* in vitro *phenotypic assays.*

Germ-free wheat plants for the Energo and Numeric varieties were produced for the several phenotypic assays described in the article as follows. Sterilized glass tubes (25cm × 5cm - 110 ml) were filled with 70 ml of autoclaved Fahraeus medium (0.132 g/L $CaCl_2$, 0.12 g/L $MgSO_4.7H_2O$, 0.1 g/L $KH_2PO_4$, 0.075 g/L $Na_2HPO_4.2H_2O$, 5 mg/L Fe-citrate, and 0.07 mg/L each of $MnCl_2.4H_2O$, $CuSO_4.5H_2O$, $ZnCl_2$, $H_3BO_3$, and $Na_2MoO_4.2H_2O$, adjusted to pH 7.5 before autoclaving) supplemented with 6 mM filtered-sterilized $CaCl_2$ (added after autoclaving). Tubes were then inclined at 30° to produce homogeneous slopes. Sterilized 3-day old seedlings obtained as described in Methods S1, were transferred in the tubes and grown on Fahraeus slopes for additional 4 days in a growth chamber at 20°C with a light/dark period of 16h/8h. Wheat roots were protected from the light by covering each tube with aluminum foil.

**Methods S5.** *Amplification of nodD gene, Illumina MiSeq sequencing and bioinformatics analysis for the quantification of Rlv strains during the co-inoculation assays.*

PCR amplifications of the nodD309 barcode sequences were performed using Phusion High-Fidelity DNA Polymerase and specific primers and conditions described in Boivin *et al*., (2020). PCR amplicons were sequenced at the GeT-PLaGE platform in Toulouse (France), using the Illumina Miseq technology with a 2×250bp paired end protocol. Paired Illumina MiSeq reads were assembled with vsearch v2.9.1 (Rognes *et al.*, 2016) using the command fastq_mergepairs and the option fastq_allowmergestagger. Demultiplexing and primer clipping were performed with cutadapt v1.9 (Martin, 2011) forcing a full-length match for sample tags and allowing a 2/3rd-length partial match for forward and reverse primers. Only reads containing both primers were retained. For each trimmed read, the expected error was estimated with vsearch's command fastq_filter and the option eeout. Each sample was then dereplicated (i.e. strictly identical reads were merged) using vsearch's command derep_fulllength, and converted to FASTA format. To prepare for clustering, samples were pooled and processed with another round of dereplication. Files containing expected error estimates were also dereplicated to retain only the lowest expected error for each unique sequence. Clustering was performed with swarm v2.1.9 (Mahé *et al.*, 2015), using a local threshold of one difference and the fastidious option. Operational taxonomic unit (OTU) representative sequences were then searched for chimeras with vsearch's command uchime_denovo, and assigned to *Rhizobium* taxa using the stampa pipeline (https://github.com/frederic-mahe/stampa/) and a custom nodD309 reference database. In parallel, each sample was individually clustered using swarm to identify local OTUs, and allow for OTUs with only one difference between their centroids. Clustering results, expected error values, taxonomic assignments and chimera detection results were used to build a raw OTU table. Up to that point, reads that could not be merged, reads without tags or primers, reads shorter than 32 nucleotides and reads with uncalled bases (“N”) had been eliminated. Additional filters were applied to keep only non-chimeric OTUs, OTUs with an expected error per nucleotide below 0.0002, OTUs containing more than 3 reads or present in at least 2 samples, and OTUs strictly identical to *Rhizobium* references (all abundant OTUs were similar to references; Table S2). After these quality controls, a total of 278382 reads for all conditions were obtained, and a mean of 11600 reads were used for each sample. In each sample, the index was calculated using the ratio between the read numbers of a given sample divided with the total read number of the sample (Table S6). Reads ratio was then converted in percentage; raw data for each sample (i.e. bacterial mixture condition and sterilization) are shown in Table S6.

**Methods S6.** *Construction of the FWPou15 GFP expressing strain.*

For the construction of FWPou15-GFP, the pHC60 (tet$^R$) plasmid (Cheng & Walker, 1998), which constitutively expresses GFP and possesses the stabilization region of the broad-host-range plasmid RK2 was introduced by electroporation in *Escherichia coli* DH5α. An ampicillin-resistant spontaneous mutant (50 mg/L Amp) of the strain FWPou15 was generated. For bacterial conjugation, 52h liquid culture of the FWPou15 Amp$^R$ strain, 24h liquid culture of *E. coli*-GFP strain and a 24h liquid culture of the *E. coli* strain carrying the pRK600 helper plasmid (Finan *et al.*, 1986) were centrifuged, washed with sterilized water and mixed at 80:50:20 ratio, respectively. A drop of each mixture was placed on TY agar plates for bacterial conjugation. Plates were incubated 48h at 24°C prior to the selection of the transformed colonies on TY agar plates containing 15 mg/L of tetracycline and 50 mg/L of ampicillin. Identity of FWPou15-GFP expressing strains was confirmed by sequencing the *nodD* gene as described above. The FWPou15-GFP expressing strain was then inoculated on wheat roots to investigate colonization by confocal microscopy (Methods S4).

**Methods S7** *Statistical analyses.*

Wheat root colonization in co-inoculation assays

To test whether significant differences occurred when *Rlv* strains were co-inoculated in wheat Energo roots, we divided the number of reads per each sample by the number of total reads of the run to obtain a relative abundance of each strain in both condition (not-sterilized roots and surface-sterilized roots). Nine values for each *strain* × *condition* obtained from 3 independent experiments were used to estimate the effect of *condition* (not-sterilized *vs* surface-sterilized roots) and the effect of the *strain*. For this, a generalized-linear-mixed model was run by using the *glmer* function implemented in lsmeans and lme4 R packages (Bates *et al.*, 2015; Lenth, 2016). Plant replicates and the temporal blocks (i.e. the 3 independent experiments) were integrated in the model as random effects. *P*-values were corrected for FDR and barplots were built using the ggplot2 and reshape2 packages (Wickham, 2007; Ginestet, 2011) on the Lsmeans obtained by correcting values for the random effects.

<u>Wheat root colonization in single strain inoculation assays</u>

To estimate the significance among the bacterial colonization abilities of *Rlv* strains in both Energo and Numeric varieties, colony forming units (cfu) obtained from all the experiments (temporal blocks, Table S6) were pooled in one unique data set as variability among replicates of each strain/condition was not significant. We first tested the effect of wheat *genotype* factor on the *cfu* factor by running a linear-mixed model with the *lmer* function by using lsmeans and lme4 R packages (Bates *et al.*, 2015; Lenth, 2016). Even the *genotype* factor was not significant, we analyzed data from both varieties independently by running a linear-mixed model where both replicates and temporal blocks were considered as random effects. *P*-values were corrected for FDR and barplots were built using the ggplot2 and reshape2 packages on the cfu Lsmeans obtained by correcting values for the random effects.

<u>Wheat root development assays</u>

The total root length and the number of later roots were analyzed on both Energo and Numeric varieties to test whether *Rlv* strains can enhance wheat root development. As we observed variability in the control plants between experiments performed with the A34 and the FWPou15 strains and those performed with the FWPou38 and IAUb11 strains, 2 data sets were analysed for these 2 classes of experiments. As experiments for A34 and FWPou15 and for FWPou38 and IAUb11 strains were made over 2 different years, it may be that not-controlled effects in the growth incubation chambers, seeds etc. could have impacted plant growth. We then compared only the experimental groups where the control plants did not show variability (Table S7). We first tested for the nested effect of *strain* × *genotype* by running a mixed-linear model as described above by including the number of replicates as random effects. As differences were observed between the 2 varieties a *lmer* model was then performed separately for each of the wheat genotypes tested. Boxplots were built using the R code with home-made scripts (Methods S8) on the raw data (not on the Lsmeans) for the total root length and the number of lateral roots.

Mycorrhiza assays

To test the impact of *Rlv* strains on *Rhizophagus irregularis* colonization, the number of AMF colonization sites when co-inoculated with each of the rhizobial strains was analyzed on both Energo and Numeric varieties. Data for the Energo variety were pooled for all the strains as they did not show variability compared to the control plants (Table S8). However, for the Numeric variety, we analyzed data on 2 sub-sets because a significant variability among the controls was observed (Table S8). The first dataset was composed of data obtained from the experiments with the A34 and the FWPou15 strains performed in parallel. The second dataset was composed of data obtained from the experiments with the FWPou38 and IAUb11 strains performed in parallel. A linear-mixed model was then run on each wheat variety independently (as the *genotype* factor explained part of the variance) with the lmer function by using lsmeans and lme4 R packages (Bates *et al.*, 2015; Lenth, 2016). The factor *condition* (presence or absence of a co-inoculated *Rlv* strain) was considered as fixed effect besides number of replicates and the temporal blocks were considered as random effects. Boxplots were built using a home-made script with the R code (Methods S8) on the raw number of colonization sites.

**Methods S8.** *R scripts used in the article.*

```
###############################################
###### Analysis on the co-inoculation assay  ##########
###############################################
library("RVAideMemoire")
library("lsmeans")
library("car")
library("vegan")
library("lme4")


data_n3=read.table("poun3.txt", h=T)
data_n6=read.table("poun6.txt", h=T)
data_n8=read.table("poun8.txt", h=T)

# Perform a GLM model on all three data set by considering the Treatment and Strain as fixed effect
F_stat <- glmer(data_n8$abd~(Treatment+Strain)^2 +(1|Rep), data = data_n8)
Anova(F_stat)
plotresid(F_stat)
lsmeans(F_stat,pairwise~(Treatment+Strain))

# Save the lsmeanes in a separate file (here called Pou_means.txt)
```

```
# Make a bar chart for each treatment mixture

library(ggplot2)
library(reshape2)

means=read.table("poun3_means.txt", h=T)


tiff("./N3.tiff", res=600, width=15, height=15, unit="cm",compression="lzw")

p<- ggplot(means, aes(x=Strain, y=lsmean, fill=Treatment)) +
geom_bar(stat="identity", color="black",
position=position_dodge()) +scale_fill_discrete(name="Treatment",
breaks=c(1, 2), labels=c("Not-sterilized", "Sterilized"))+
xlab("Strains")+ylab("Mean abundance")+
coord_flip() +
scale_fill_brewer(palette = "Set3")

# print(p) # Use it only to visualized the graph before save it

# Personalized the plot


p+labs(title="a", x="Strains", y = "Mean abundance")+
theme_classic() +
scale_fill_brewer(palette = "Greys")

dev.off()


# Second mixture

means2= read.table("poun6_means.txt", h=T)


tiff("./N6.tiff", res=600, width=15, height=15, unit="cm",compression="lzw")


p2<- ggplot(means2, aes(x=Strain, y=lsmean, fill=Treatment)) +
geom_bar(stat="identity", color="black",
position=position_dodge()) +
scale_fill_discrete(name="Treatment",
breaks=c(1, 2),
labels=c("Not-sterilized", "Sterilized"))+
xlab("Strains")+ylab("Mean abundance")+
coord_flip() +
scale_fill_brewer(palette = "Set3")


# print(p2) # use it only to visualized the graph before save it

# Personalized the plot

p2+labs(title="b", x="Strains", y = "Mean abundance")+
theme_classic() +
scale_fill_brewer(palette = "Greys")

dev.off()


# Third mixture

means3=read.table("poun8_means.txt", h=T)


tiff("./N8.tiff", res=600, width=15, height=15, unit="cm",compression="lzw")
p3<- ggplot(means3, aes(x=Strain, y=lsmean, fill=Treatment)) +
```

```
geom_bar(stat="identity", color="black",
position=position_dodge()) +
scale_fill_discrete(name="Treatment",
breaks=c(1, 2),
labels=c("Not-sterilized", "Sterilized"))+
xlab("Strains")+ylab("Mean abundance")+
coord_flip() +
scale_fill_brewer(palette = "Set3")

# print(p3) # use it only to visualized the graph before save it
# Personalized the plot

p3+labs(title="c", x="Strains", y = "Mean abundance")+
theme_classic() +
scale_fill_brewer(palette = "Greys")

dev.off()

##########################################################
########### Analysis on the bacterial root colonization assay ########
##########################################################

setwd("R:/CGJC_2016_17-18/wheat-project/RHIZOWHEAT_project/Phenotypic test/Growth-curves")

library("RVAideMemoire")
library("lsmeans")
library("car")
library("vegan")
library("lme4")

dataG=read.table("growth_stat21022020.txt", h=T)
# Run a model with all factors
F_stat <- lmer(dataG$cfu~(treatment+variety+strain)^2 +(1|block/rep), data=dataG)
Anova(F_stat)

##### Run analysis per each variety
## Energo

dataEne=dataG [which(dataG$variety=='Energo'),]

F_stat <- lmer(dataEne$cfu~(treatment+strain)^2 +(1|block/rep), data=dataEne)
Anova(F_stat)
lsmeans(F_stat,pairwise~(strain+treatment))

# Make a plot for Energo

means_Ene=read.table("energo_lsmeans.txt", h=T)

library(ggplot2)
tiff("./bacterial_growth_energo.tiff", res=600, width=15, height=15, unit="cm",compression="lzw")
```

```
p<- ggplot(means_Ene, aes(x=strain, y=lsmean, fill=treatment)) +
geom_bar(stat="identity", color="black",
position=position_dodge()) +
geom_errorbar(aes(ymin=lsmean-SE, ymax=lsmean+SE), width=.2,
position=position_dodge(.9))

# print(p) # use it only to visualized the graph before save it


# Personalized the plot

p+labs(title="Bacterial colonization (Energo)", x="Strains", y = "cfu/mg(log10)")+
theme_classic() +
scale_fill_brewer(palette = "Greys")


dev.off()

# Numéric

dataNum=dataG [which(dataG$variety=='Numeric'),]

F_stat <- lmer(dataNum$cfu~(treatment+strain)^2 +(1|block/rep), data=dataNum)
Anova(F_stat)
lsmeans(F_stat,pairwise~(strain+treatment))

## Make a plot for numéric

means_Num=read.table("numeric_lsmeans.txt", h=T)

tiff("./bacterial_growth_numeric.tiff", res=600, width=15, height=15, unit="cm",compression="lzw")


p<- ggplot(means_Num, aes(x=strain, y=lsmean, fill=treatment)) +
geom_bar(stat="identity", color="black",
position=position_dodge()) +
geom_errorbar(aes(ymin=lsmean-SE, ymax=lsmean+SE), width=.2,
position=position_dodge(.9))

# print(p) # use it only to visualized the graph before save it

# Personalized the plot

p+labs(title="Bacterial colonization (Numeric)", x="Strains", y = "cfu/mg(log10)")+
theme_classic() +
scale_fill_brewer(palette = "Greys")


dev.off()


##########################################################

########### Analysis on wheat root development ###############

##########################################################


setwd('R:/CGJC_2016_17-18/wheat-project/RHIZOWHEAT_project/Phenotypic test/Later root')


library("RVAideMemoire")
library("lsmeans")
library("car")
library("vegan")
library("lme4")
```

```
############################################
####### Analysis on FWPou15 and A34 strains#######
############################################

data1= read.table('Roots_Pou15_A34.txt', header=T, sep="\t")

# linear model on both varieties (lenght)

F_stat <- lmer(data1$lenght~(strain+variety)^2 + (1|rep), data=data1)
Anova(F_stat)
# on the Energo variety

dataEne=data1 [which(data1$variety=='energo'),]
F_stat <- lmer(dataEne$laterl.roots~(strain)^2 + (1|rep), data=dataEne)
Anova(F_stat)
lsmeans(F_stat,pairwise~(strain))

# On the Numéric variety

dataNum=data1 [which(data1$variety=='numeric'),]
F_stat <- lmer(dataNum$laterl.roots~(strain)^2 + (1|rep), data=dataNum)
Anova(F_stat)
lsmeans(F_stat,pairwise~(strain))

#### Make the boxplot for all dataset
# Total root length

tiff("./root-develp_1.tiff", res=600, width=15, height=15, unit="cm",compression="lzw")

boxplot(lenght~strain*variety, data=data1, xlab="", ylab = "Total root lenght", main = "", cex.axis=0.7,las=2,
at = c(1,2,3, 5,6,7), axes=F)
box()
axis(2, at=seq(0,140,20), seq(0,140,20), las=1)
axis(1, at=c(1,2,3, 5,6,7),labels = c("A34","ctr","FWPou15", "A34","ctr","FWPou15"), cex.axis=0.7,las=2,side=1)
mtext("Energo", side = 1, line = 3.8, adj = 0.250, cex=0.7, font=2)
mtext("Numéric", side = 1, line = 3.8, adj = 0.830, cex=0.7, font=2)
axis(1, at=c(0.5,1,2,3.5), line=3.8, tick=T, labels=rep("",4), lwd=1, lwd.ticks=0)
axis(1, at=4+c(0.5,1,2,3.5), line=3.8, tick=T, labels=rep("",4), lwd=1, lwd.ticks=0)

dev.off()

# Lateral roots

tiff("./lateral_roots_1.tiff", res=600, width=15, height=15, unit="cm",compression="lzw")

boxplot(laterl.roots~strain*variety, data=data1, xlab="", ylab = "N° lateral roots", main = "", cex.axis=0.7,las=2,
at = c(1,2,3, 5,6,7), axes=F)
box()
axis(2, at=seq(0,120,20), seq(0,120,20), las=1)
axis(1, at=c(1,2,3, 5,6,7),labels = c("A34","ctr","FWPou15", "A34","ctr","FWPou15"), cex.axis=0.7,las=2,side=1)
mtext("Energo", side = 1, line = 3.8, adj = 0.250, cex=0.7, font=2)
```

```
mtext("Numéric", side = 1, line = 3.8, adj = 0.830, cex=0.7, font=2)
axis(1, at=c(0.5,1,2,3.5), line=3.8, tick=T, labels=rep("",4), lwd=1, lwd.ticks=0)
axis(1, at=4+c(0.5,1,2,3.5), line=3.8, tick=T, labels=rep("",4), lwd=1, lwd.ticks=0)

dev.off()

##############################################
######## Analysis on FWPou15 and A34 strains######
##############################################

data2=read.table("Roots_IAUb_Pou38.txt",h=T, sep="\t")


# Linear model on both varieties


F_stat <- lmer(data2$lenght~(condition+variety)^2 + (1|rep), data=data2)
Anova(F_stat)
F_stat <- lmer(data1$lateral.roots~(condition+variety)^2 + (1|rep), data=data1)
Anova(F_stat)


# On the Energo variety


dataEne=data2 [which(data2$variety=='energo'),]
F_stat <- lmer(dataEne$lenght~(condition)^2 + (1|rep), data=dataEne)
Anova(F_stat)
lsmeans(F_stat,pairwise~(condition))


# On the Numéric variety


dataNum=data2 [which(data2$variety=='numeric'),]
F_stat <- lmer(dataNum$lateral.roots~(condition)^2 + (1|rep), data=dataNum)
Anova(F_stat)
lsmeans(F_stat,pairwise~(condition))


################################################
######### Make a unique boxplot for all dataset#######
################################################


tiff("./root-develp_2.tiff", res=600, width=15, height=15, unit="cm",compression="lzw")

boxplot(lenght~condition*variety, data=data2, xlab="", ylab = "Total root lenght", main = "", cex.axis=0.7,las=2,
at = c(1,2,3, 5,6,7), axes=F)
box()
axis(2, at=seq(0,350,50), seq(0,350,50), las=1)
axis(1, at=c(1,2,3, 5,6,7),labels = c("ctr","FWPou38","IAUb11", "ctr","FWPou38","IAUB11"), cex.axis=0.7,las=2,side=1)
mtext("Energo", side = 1, line = 3.8, adj = 0.250, cex=0.7, font=2)
mtext("Numéric", side = 1, line = 3.8, adj = 0.830, cex=0.7, font=2)
axis(1, at=c(0.5,1,2,3.5), line=3.8, tick=T, labels=rep("",4), lwd=1, lwd.ticks=0)
axis(1, at=4+c(0.5,1,2,3.5), line=3.8, tick=T, labels=rep("",4), lwd=1, lwd.ticks=0)

dev.off()


# Lateral roots
```

```
tiff("./lateral_roots_2.tiff", res=600, width=15, height=15, unit="cm",compression="lzw")

boxplot(lateral.roots~condition*variety, data=data2, xlab="", ylab = "N° lateral roots", main = "", cex.axis=0.7,las=2,
at = c(1,2,3, 5,6,7), axes=F)
box()
axis(2, at=seq(0,200,50), seq(0,200,50), las=1)
axis(1, at=c(1,2,3, 5,6,7),labels = c("ctr","FWPou38","IAUb11", "ctr","FWPou38","IAUb11"), cex.axis=0.7,las=2,side=1)
mtext("Energo", side = 1, line = 3.8, adj = 0.250, cex=0.7, font=2)
mtext("Numéric", side = 1, line = 3.8, adj = 0.830, cex=0.7, font=2)
axis(1, at=c(0.5,1,2,3.5), line=3.8, tick=T, labels=rep("",4), lwd=1, lwd.ticks=0)
axis(1, at=4+c(0.5,1,2,3.5), line=3.8, tick=T, labels=rep("",4), lwd=1, lwd.ticks=0)

dev.off()


### Put the 4 box plots in the same image


tiff("./root-develp_all.tiff", res=600, width=15, height=15, unit="cm",compression="lzw")

par(mfrow=c(2,2))
P1= boxplot(lenght~condition*variety, data=data1, xlab="", ylab = " Total root lenght ", main = "a", cex.axis=0.7,las=2,
at = c(1,2,3, 5,6,7), axes=F)
box()
axis(2, at=seq(0,140,20), seq(0,140,20), las=1)
axis(1, at=c(1,2,3, 5,6,7),labels = c("ctr","A34","FWPou15", "ctr","A34","FWPou15"), cex.axis=0.7,las=2,side=1)
mtext("Energo", side = 1, line = 3.8, adj = 0.250, cex=0.7, font=2)
mtext("Numeric", side = 1, line = 3.8, adj = 0.830, cex=0.7, font=2)
axis(1, at=c(0.5,1,2,3.5), line=3.8, tick=T, labels=rep("",4), lwd=1, lwd.ticks=0)
axis(1, at=4+c(0.5,1,2,3.5), line=3.8, tick=T, labels=rep("",4), lwd=1, lwd.ticks=0)


P2= boxplot(laterl.roots~condition*variety, data=data1, xlab="", ylab = "N° lateral roots", main = "b", cex.axis=0.7,las=2,
at = c(1,2,3, 5,6,7), axes=F, main="b")
box()
axis(2, at=seq(0,120,20), seq(0,120,20), las=1)
axis(1, at=c(1,2,3, 5,6,7),labels = c("ctr","A34","FWPou15", "ctr","A34","FWPou15"), cex.axis=0.7,las=2,side=1)
mtext("Energo", side = 1, line = 3.8, adj = 0.250, cex=0.7, font=2)
mtext("Numeric", side = 1, line = 3.8, adj = 0.830, cex=0.7, font=2)
axis(1, at=c(0.5,1,2,3.5), line=3.8, tick=T, labels=rep("",4), lwd=1, lwd.ticks=0)
axis(1, at=4+c(0.5,1,2,3.5), line=3.8, tick=T, labels=rep("",4), lwd=1, lwd.ticks=0)

P3=boxplot(lenght~condition*variety, data=data2, xlab="", ylab = " Total root lenght ", main = "c", cex.axis=0.7,las=2,
at = c(1,2,3, 5,6,7), axes=F, main="c")
box()
axis(2, at=seq(0,350,50), seq(0,350,50), las=1)
axis(1, at=c(1,2,3, 5,6,7),labels = c("ctr","FWPou38","IAUb11", "ctr","FWPou38","IAUB11"), cex.axis=0.7,las=2,side=1)
mtext("Energo", side = 1, line = 3.8, adj = 0.250, cex=0.7, font=2)
mtext("Numeric", side = 1, line = 3.8, adj = 0.830, cex=0.7, font=2)
axis(1, at=c(0.5,1,2,3.5), line=3.8, tick=T, labels=rep("",4), lwd=1, lwd.ticks=0)
axis(1, at=4+c(0.5,1,2,3.5), line=3.8, tick=T, labels=rep("",4), lwd=1, lwd.ticks=0)

P4= boxplot(lateral.roots~condition*variety, data=data2, xlab="", ylab = "N° lateral roots", main = "d", cex.axis=0.7,las=2,
at = c(1,2,3, 5,6,7), axes=F, main="d")
box()
axis(2, at=seq(0,200,50), seq(0,200,50), las=1)
axis(1, at=c(1,2,3, 5,6,7),labels = c("ctr","FWPou38","IAUb11", "ctr","FWPou38","IAUb11"), cex.axis=0.7,las=2,side=1)
mtext("Energo", side = 1, line = 3.8, adj = 0.250, cex=0.7, font=2)
mtext("Numeric", side = 1, line = 3.8, adj = 0.830, cex=0.7, font=2)
axis(1, at=c(0.5,1,2,3.5), line=3.8, tick=T, labels=rep("",4), lwd=1, lwd.ticks=0)
axis(1, at=4+c(0.5,1,2,3.5), line=3.8, tick=T, labels=rep("",4), lwd=1, lwd.ticks=0)

dev.off()
```

```
#####################################
##### Analysis on AMF colonization#########
##### in presence of Rlv strains ############
######################################

setwd ("R:/CGJC_2016_17-18/wheat-project/RHIZOWHEAT_project/Phenotypic test/AMF-colonisation")

# Boxplot for whole dataset Energo+ Numéric

data=read.table("Statistics_all.txt", h=T, sep="\t")

tiff("./AMF-colonization-all.tiff", res=600, width=15, height=15, unit="cm",compression="lzw")

boxplot(sites~condition*variety, data=data, xlab="", ylab = "Colonization sites", main = "", cex.axis=0.7,las=2,
at = c(1,2,3,4,5, 7,8,9,10,11), axes=F)
box()
axis(2, at=seq(0,200,50), seq(0,200,50), las=1)
axis(1, at=c(1,2,3,4,5, 7,8,9,10,11),labels = c("AMF","A34","FWPou15","FWPou38","IAUb11","AMF","A34","FWPou15", "",
""), cex.axis=0.7,las=2,side=1)
mtext("Energo", side = 1, line = 3.8, adj = 0.250, cex=0.7, font=2)
mtext("Numéric", side = 1, line = 3.8, adj = 0.830, cex=0.7, font=2)
axis(1, at=c(0.5,1,2,5.5), line=3.8, tick=T, labels=rep("",4), lwd=1, lwd.ticks=0)
axis(1, at=6+c(0.5,1,2,5.5), line=3.8, tick=T, labels=rep("",4), lwd=1, lwd.ticks=0)

dev.off()

# Plot on second experiment numéric


data2= read.table("numeric_second.txt", h=T, sep="\t")

tiff("./AMF-colonization-numeric.tiff", res=600, width=15, height=15, unit="cm",compression="lzw")

boxplot(sites~condition*variety, data=data2, xlab="", ylab = "Colonization sites", main = "", cex.axis=0.7,las=2,
at = c(1,2,3), axes=F)
box()
axis(2, at=seq(0,250,50), seq(0,250,50), las=1)
axis(1, at=c(1,2,3),labels = c("AMF","FWPou38","AUb11"), cex.axis=0.7,las=2,side=1)
mtext("Numéric", side = 1, line = 3.8, adj = 0.450, cex=0.7, font=2)
axis(1, at=c(0.5,1,2,5.5), line=3.8, tick=T, labels=rep("",4), lwd=1, lwd.ticks=0)

dev.off()

# Combine figures


data1= read.table("energo1.txt", h=T, sep="\t")
data2= read.table("energo2.txt", h=T, sep="\t")
data3= read.table("numeric1.txt", h=T, sep="\t")
data4= read.table("numeric2.txt", h=T, sep="\t")

tiff("./AMFColonization_all.tiff", res=600, width=15, height=15, unit="cm",compression="lzw")

par(mfrow=c(2,2))

P1= boxplot(sites~condition*variety, data=data1, xlab="", ylab = "Colonization sites", main = "a", cex.axis=0.7,las=2,
at = c(1,2,3), axes=F)
box()
axis(2, at=seq(0,200,50), seq(0,200,50), las=1)
```

```
axis(1, at=c(1,2,3),labels = c("AMF","A34","FWPou15"), cex.axis=0.7,las=2,side=1)
mtext("Energo", side = 1, line = 3.8, adj = 0.450, cex=0.7, font=2)
axis(1, at=c(0.5,1,2,5.5), line=3.8, tick=T, labels=rep("",4), lwd=1, lwd.ticks=0)

P2=boxplot(sites~condition*variety, data=data2, xlab="", ylab = "Colonization sites", main = "b", cex.axis=0.7,las=2,
at = c(1,2,3), axes=F)
box()
axis(2, at=seq(0,100,20), seq(0,100,20), las=1)
axis(1, at=c(1,2,3),labels = c("AMF","FWPou38","IAUb11"), cex.axis=0.7,las=2,side=1)
mtext("Energo", side = 1, line = 3.8, adj = 0.450, cex=0.7, font=2)
axis(1, at=c(0.5,1,2,5.5), line=3.8, tick=T, labels=rep("",4), lwd=1, lwd.ticks=0)

P3=boxplot(sites~condition*variety, data=data3, xlab="", ylab = "Colonization sites", main = "c", cex.axis=0.7,las=2,
at = c(1,2,3), axes=F)
box()
axis(2, at=seq(0,150,50), seq(0,150,50), las=1)
axis(1, at=c(1,2,3),labels = c("AMF","A34","FwPou15"), cex.axis=0.7,las=2,side=1)
mtext("Numeric", side = 1, line = 3.8, adj = 0.450, cex=0.7, font=2)
axis(1, at=c(0.5,1,2,5.5), line=3.8, tick=T, labels=rep("",4), lwd=1, lwd.ticks=0)

P4=boxplot(sites~condition*variety, data=data4, xlab="", ylab = "Colonization sites", main = "d", cex.axis=0.7,las=2,
at = c(1,2,3), axes=F)
box()
axis(2, at=seq(50,250,50), seq(50,250,50), las=1)
axis(1, at=c(1,2,3),labels = c("AMF","FWPou38","IAUb11"), cex.axis=0.7,las=2,side=1)
mtext("Numeric", side = 1, line = 3.8, adj = 0.450, cex=0.7, font=2)
axis(1, at=c(0.5,1,2,5.5), line=3.8, tick=T, labels=rep("",4), lwd=1, lwd.ticks=0)

dev.off()


# Linear model per variery

# Energo


data=read.table("Statistics_all.txt", h=T, sep="\t")

dataEn=data [which(data$variety=='Energo'),]

AMF_F_stat <- lmer(dataEn$sites~(condition)^2 +(1|rep/block), data = dataEn)
Anova(AMF_F_stat)
plotresid(AMF_F_stat)
lsmeans(AMF_F_stat,pairwise~(condition))

dataNum=data [which(data$variety=='Numeric'),]

AMF_F_stat <- lmer(dataNum$sites~(condition)^2 +(1|rep/block), data = dataNum)
Anova(AMF_F_stat)
plotresid(AMF_F_stat)
lsmeans(AMF_F_stat,pairwise~(condition))

data2=read.table("numeric_second.txt", h=T, sep="\t")

AMF_F_stat <- lmer(data2$sites~(condition)^2 +(1|rep/block), data = data2)
Anova(AMF_F_stat)
plotresid(AMF_F_stat)
lsmeans(AMF_F_stat,pairwise~(condition))
```

**Methods S9**

**Supplemental references**

**Bates D, Mächler M, Bolker BM, Walker SC**. **2015**. Fitting linear mixed-effects models using lme4. *Journal of Statistical Software* **67**: 1–48.

**Boivin S, Mahé F, Prevent M, Tancelin M, Tauzin Ma, Wielbo J, Mazurier S, Young P, Lepetit M**. **2020**. Genetic variation in host-specific competitiveness of *Rhizobium leguminosarum* symbiovar *viciae* . *Authorea* doi: 10.22541/au.159237007.72934061.

**Cheng HP, Walker GC**. **1998**. Succinoglycan is required for initiation and elongation of infection threads during nodulation of alfalfa by *Rhizobium meliloti*. *Journal of Bacteriology* **180**: 5183–5191.

**Finan TM, Kunkel B, De Vos GF, Signer ER**. **1986**. Second symbiotic megaplasmid in *Rhizobium meliloti* carrying exopolysaccharide and thiamine synthesis genes. *Journal of Bacteriology* **167**: 66–72.

**Wickham H**. **2016**. ggplot2: Elegant Graphics for Data Analysis. Springer-Verlag New York. ISBN 978-3-319-24277-4, https://ggplot2.tidyverse.org.

**Kolde R, Vilo J**. **2015**. GOsummaries: An R Package for Visual Functional Annotation of Experimental Data **4**: 574.

**Lenth R V. 2016**. Least-squares means: The R package lsmeans. *Journal of Statistical Software* **69**: 1–33.

**Mahé F, Rognes T, Quince C, de Vargas C, Dunthorn M**. **2015**. Swarmv2: Highly-scalable and high-resolution amplicon clustering. *PeerJ* **3**: e1420.

**Martin M**. **2011**. Cutadapt removes adapter sequences from high-throughput sequencing reads. *EMBnet.journal*. doi: https://doi.org/10.14806/ej.17.1.200.

**Rognes T, Flouri T, Nichols B, Quince C, Mahé F**. **2016**. VSEARCH: A versatile open source tool for metagenomics. *PeerJ* **4**:e2584

**Sallet E, Gouzy J, Schiex T**. **2014**. EuGene-PP: A next-generation automated annotation pipeline for prokaryotic genomes. *Bioinformatics* **30**: 2659–2661.

**Tamura K, Peterson D, Peterson N, Stecher G, Nei M, Kumar S**. **2011**. MEGA5: Molecular evolutionary genetics analysis using maximum likelihood, evolutionary distance, and maximum parsimony methods. *Molecular Biology and Evolution* **25**: 2731–2739.

**Wickham H**. **2007**. Reshaping data with the reshape package. *Journal of Statistical Software* **21**: 1–20.

**Xia X**. **2017**. DAMBE6: New tools for microbial genomics, phylogenetics, and molecular evolution. *Journal of Heredity*. **108**: 431–437.
